# Growth-adaptive spring electronics for long-term, same-neuron mapping in the developing rat brain

**DOI:** 10.64898/2026.02.16.705931

**Authors:** Ariel J. Lee, Hao Sheng, Arnau Marin-Llobet, Zheliang Wang, Jaeyong Lee, Ren Liu, Xinhe Zhang, Emma Hsiao, Jongmin Baek, Almir Aljovic, Ding Liu, Yichun He, Nanshu Lu, Jia Liu

## Abstract

Neural activity reorganizes profoundly after birth, transitioning from highly synchronous population events to sparse, decorrelated firing in the mature brain. Although inhibitory maturation and shifts in excitation-inhibition balance have been implicated in this process, how individual neurons implement the transition remains unclear because rapid brain growth has prevented long-term, same-neuron mapping. Here, we introduce growth-adaptive spring electronics that provide depth-wise compliance during tissue expansion, maintaining a stable electrode-tissue interface over weeks of neonatal development. We developed a vision-language model-assisted spike processing pipeline for the developing brain that probabilistically matches units across days using high-density waveform spatial footprints, despite developmental changes in the neonatal brain. Together, these innovations enable spike-resolved mapping of the same neurons in rat visual cortex and medial prefrontal cortex from postnatal day 10 to 45. Using population coupling to quantify each neuron’s coordination with local population activity, we show that developmental decorrelation is driven primarily by a distinct subset of neurons that progressively shifts from strong to weak coupling during postnatal weeks 3 to 5, whereas other neurons remain stably weakly or strongly coupled throughout development. These results resolve population-level desynchronization into identifiable neuron-specific trajectories. This framework enables direct tests in neurodevelopmental disorder models, including schizophrenia and autism, of whether altered maturation reflects global circuit imbalance or selective disruption and mistiming of specific developmental programs.

## Main text

Understanding how neuronal activity reorganizes during early postnatal development requires mapping activity in the same animal over time with single-cell, single-spike spatiotemporal resolution. Shortly after birth, the mammalian brain undergoes rapid maturation accompanied by significant tissue expansion and displacement, while neural circuits transition from highly synchronous activity to increasingly sparse and decorrelated activity patterns. These changes are thought to support efficient information processing and the emergence of mature circuit function^1–6^. However, current recording technologies have made continuous, single-cell-resolved electrical mapping across this dynamic period difficult. Consequently, the cellular mechanisms underlying these network-level transitions have been inferred from cross-sectional recordings collected from different animals at different developmental stages, rather than observed longitudinally from the same cells within the same individual animals.

In addition, existing recording modalities do not fully address this gap. Functional magnetic resonance imaging and related methods enable non-invasive continuous monitoring of brain activity, but lack the spatiotemporal resolution needed to resolve single-neuron activity^3,7,8^. Optical imaging with calcium or voltage indicators can record cellular activity in early development, but remains constrained by temporal resolution, imaging depth, and signal instability (e.g. photobleaching), as well as by the difficulty of maintaining stable cranial windows and reliably re-registering the same field of view across weeks of skull growth and tissue displacement^9–12^. Recent two-photon imaging studies have tracked activity from the same neurons for days during early postnatal stages and reported progressive decorrelation as inhibitory circuits mature, but these approaches remain limited in duration and do not provide continuous, single-unit recordings across early postnatal stages^13,14^. Long-term intracortical recordings in the rapidly developing brain are constrained by mechanical challenges. The postnatal brain expands and shifts relative to the skull and implanted devices, whereas conventional rigid silicon or semi-flexible polymer probes are orders of magnitude stiffer than brain tissue^15,16^. This mechanical mismatch promotes relative motion at the electrode-tissue interface, leading to signal drift and preventing stable recording from the same local tissue volume over time. It also induces inflammation, glial encapsulation, and neuronal loss near the implants, which degrades recording quality even in adult systems^17–19^ and may be more severe during early postnatal development.

Advances in flexible electronics have reduced mechanical mismatch and improved chronic stability in adult brain recordings. By mimicking the neuron-level size and mechanical properties, tissue-like electronics can minimize micromotion-induced damage and support stable single-unit recordings over months^20,21^. More recently, tissue-like stretchable nanoelectronics have been integrated into developing tissues, accommodating two-dimensional (2D) cell layers as they fold into 3D organoids *in vitro*^22–24^ and enabling integration during embryonic development *in vivo*^25^. These tissue-like electronics designs show the potential of tissue-growth-compatible bioelectronics, but they have not yet enabled continuous recordings in the intact, rapidly developing mammalian brain *in vivo*, where tissue expansion and displacement must be accommodated.

To address these challenges, here, we introduce a growth-adaptive intracortical interface engineered specifically to deform along with the growing rat brain (Fig. 1a). The device is fabricated as a planar spiral using standard microfabrication processes and, upon implantation, transforms into a 3D helical spring embedded within the tissue. This helical geometry allows for high compliance along the primary axis of postnatal cortical expansion while maintaining local electrode-tissue alignment. Combined with an analysis pipeline that adapts conventional spike sorting for the developing brain, this platform enables continuous mapping of identified single units from the neonatal period into adulthood despite large changes in brain size and intrinsic cellular properties.

**Fig. 1.**
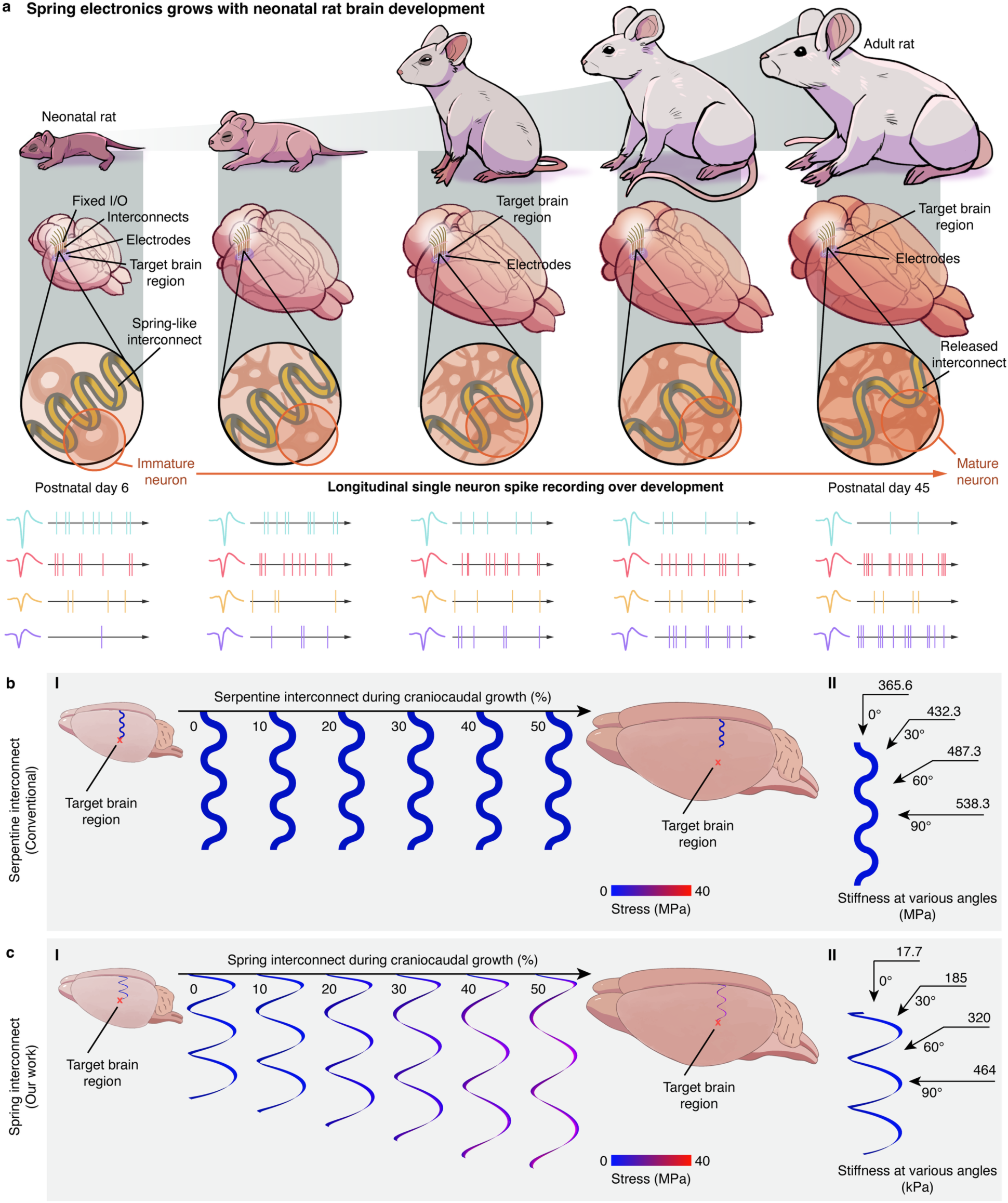
Spring electronics that accommodate brain growth for longitudinal single-neuron activity mapping in neonatal rats. **a**, Conceptual illustration of a stretchable, spring-like electrode array implanted into the neonatal rat brain. As the brain grows, the interconnect deforms with the surrounding tissue, enabling stable mapping of single-neuron dynamics and revealing development-dependent single-neuron activity changes. **b,c**, (I) Finite element analysis (FEA) of von Mises stress (MPa) in a representative serpentine **(b)** versus a spring **(c)** interconnect under 0%, 10%, 20%, 30%, 40% and 50% expansion along the craniocaudal axis. Schematics compares a serpentine **(b)** versus a spring **(c)** interconnect embedded in the brain tissue before and after growth, with one end anchored to the skull. (II) Effective stiffness of the representative serpentine **(b)** and spring **(c)** interconnect at out-of-plane stretching angles of 0°, 30°, 60° and 90°.

We apply these growth-adaptive electronics to record from visual cortex (V1) and medial prefrontal cortex (mPFC) in rats from postnatal day 10 (P10) to P45, spanning the period in which cortical activity decorrelates and behavioral function matures. Same-neuron mapping during development shows that in these circuits, development-associated cortical activity decorrelation arises predominantly from a subset of neurons that change their relationship to the local population, rather than a uniform adjustment across all neurons, which could not be observed through the traditional cross-sectional view of the brain dynamics.

### Designing stretchable electronics for neonatal brain

We previously developed stretchable electronics that can be implanted into organoids and aquatic animal embryos throughout the tissue development, enabling longitudinal tracking of cellular activity during development^22,24,25^. However, these techniques are challenging to apply to mammalian brain development studies, which typically require implantation of a stretchable device into an already-established neural network. This imposes two additional design constraints. First, the implanted electronics must accommodate postnatal brain growth so that recording sites remain aligned with the same brain micro-region at single-cell precision. Second, the device must preserve its stretchable, deformable architecture after implantation. The rapidly developing mammalian brain therefore presents a unique mechanical challenge for long-term stable electrical interfacing.

In standard stereotaxic implantation, devices are inserted to a defined depth, while the input/output (I/O) interface is rigidly anchored to the skull^26–28^ (Extended Data Fig. 1a).

Therefore, as a first step, we considered a simplified model in which a stretchable device is already embedded in the neonatal brain and its I/O end is anchored at the skull. We then asked whether such a structure could elongate sufficiently during development, especially along the depth axis, to maintain alignment between electrodes and their targeted neurons.

We first examined a conventional serpentine structure, a standard architecture for stretchable electronics that we previously used in organoids and embryos^22,24,25^. Using finite element analysis (FEA), we modeled a conventional serpentine interconnect comprising 3/50/3-nm chromium/gold (Cr/Au/Cr) conductive layers encapsulated by a 1-µm-thick rigid polymer (e.g., SU-8) layer (Fig. 1b, I, Methods). Under 50% craniocaudal tissue expansion, this structure showed negligible elongation (0.184%). We embedded serpentine interconnects inside a hydrogel brain phantom to experimentally confirm this result. We monitored deformation of the serpentine structure as the hydrogel swelled. However, the elongation of the serpentine did not occur (Extended Data Fig. 1b). We then systematically optimized serpentine geometry to increase stretchability. However, even the most compliant designs achieved <6% elongation under 30% expansion (Extended Data Fig. 1c). Replacing the rigid SU-8 with perfluoropolyether-dimethacrylate (PFPE-DMA), an ultrasoft polymer previously used in embryonic systems^25^, improved elongation but remained insufficient relative to developmental brain expansion (1.3% under 30% craniocaudal expansion, Extended Data Fig. 1d).

This limitation reflects a fundamental mismatch between serpentine design assumptions and the mechanical environment of the developing brain. Serpentines are primarily optimized for in-plane uniaxial stretching, whereas postnatal brain growth imposes nearly isotropic tensile stress.

Achieving preferential elongation along a single axis under isotropic loading requires strong mechanical anisotropy, a substantially lower axial stiffness along that elongation axis than along orthogonal directions. Conventional serpentine geometries, however, provide limited anisotropy and exhibit comparable effective axial stiffness across axes (Fig. 1b, II).

In contrast, from a mechanical engineering perspective, a helical spring structure offers precisely this anisotropy: axial extension is converted into bending and torsion of the coils, yielding very low effective stiffness along the spring axis while maintaining higher stiffness laterally, thereby promoting deformation along the spring axis even under isotropic loading. A complementary intuition comes from the ribbon’s cross-section imposed by microfabrication techniques. Due to microfabrication constraints, the interconnect ribbons have a highly anisotropic cross-section, with thickness *t* of several µm and width *w* of tens to hundreds of µm (*w* ≫ *t*). In planar serpentine interconnects, in-plane stretching is accommodated primarily by bending about the out-of-plane axis (normal to the fabrication plane) of the rectangular ribbon cross-section, for which the flexural rigidity is proportional to *tw*^3^/12. In the helical spring interconnect, axial extension is accommodated mainly by ribbon out-of-plane bending and torsion, bending about the in-plane width axis, for which flexural rigidity becomes proportional to *wt*^3^/12. Because *w* ≫ *t*, the latter is smaller by a factor of (*t*/*w*)^2^, providing an additional source of effective axial softness along the spring axis.

Guided by this principle, we designed a spring-like interconnect to accommodate depth-wise elongation in an isotropically expanding neonatal brain (Fig. 1c, I). FEA using the same material stack and comparable dimensions as the previous serpentine designs showed that the spring structure deforms synchronously with tissue growth, effectively matching depth-wise elongation when embedded in brain tissue. Simulations further showed a marked reduction in effective stretch stiffness along the longitudinal axis (Fig. 1c, II). Together, these results suggest that compared to serpentine structure, helical spring-like structure can accommodate early neonatal brain development more seamlessly (Extended Data Fig. 1e).

### Spiral-to-spring transformation via implantation

Implantation of a helical spring-like device in the neonatal brain, however, poses two challenges. First, single-unit isolation requires high-density microelectrode arrays, which are typically microfabricated on planar wafers. Second, conventional stereotaxic implantation can cause highly stretchable structures to collapse, buckle, or stiffen during implantation, undermining the very deformability needed for brain development accommodation. Together, these constraints create a fabrication and implantation paradox: helical springs provide the required mechanical anisotropy but are not directly compatible with planar lithography and standard implantation workflows.

To resolve this, we developed a spiral-to-spring transformation strategy. Devices are fabricated as planar spirals using standard microfabrication, then convert into 3D helical springs in situ during stereotaxic implantation (Fig. 2a). This approach preserves photolithographic precision for microelectronics fabrication, while achieving the desired spring structure after implantation through a simple surgical procedure.

**Fig. 2.**
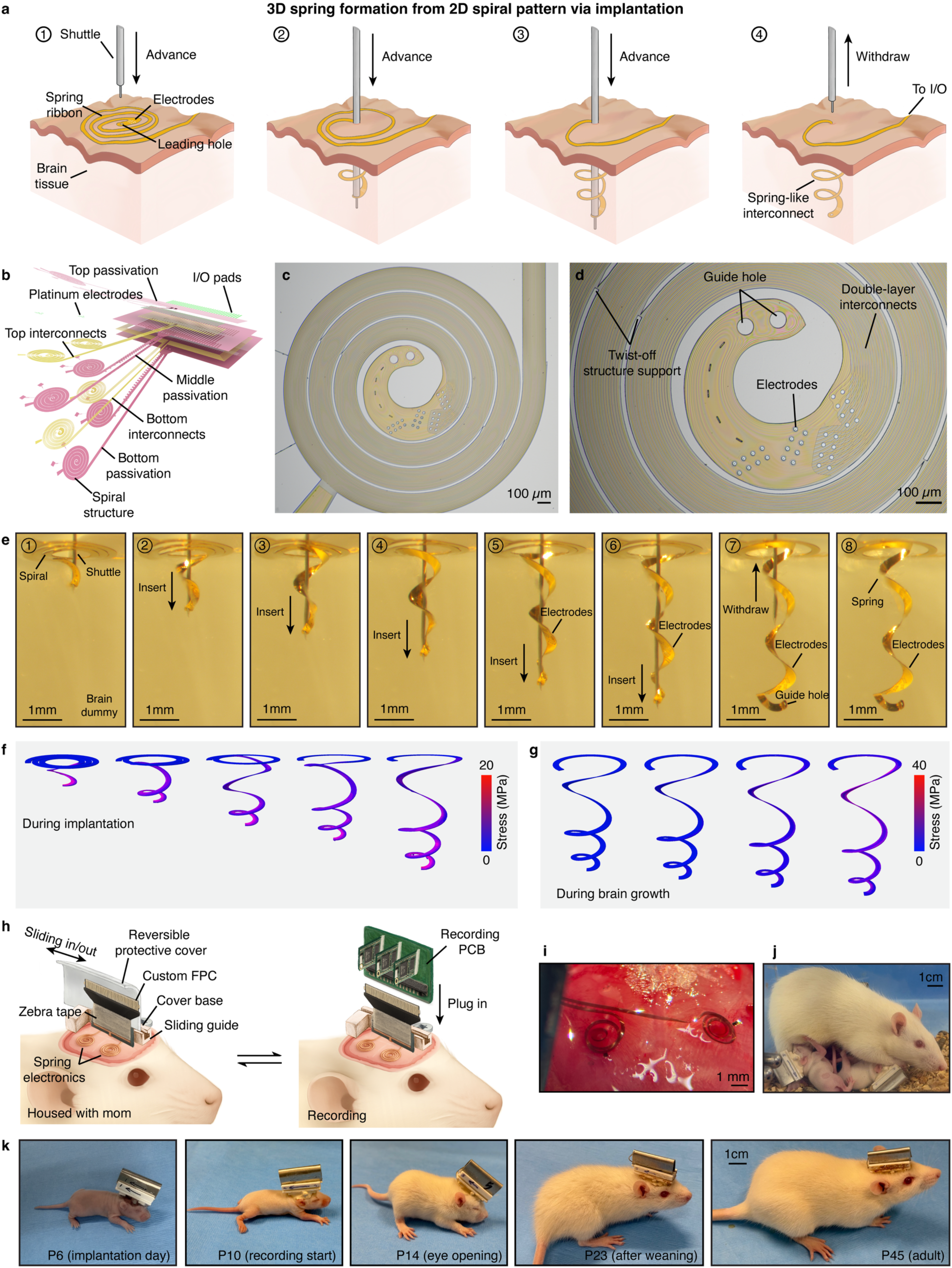
Implantation-induced formation of a spring deformable brain implant. **a**, Schematics of the planar spiral device and stepwise implantation that converts it into a spring-like structure in the brain tissue. (1) A planar flexible spiral device is placed on the brain surface, with a guide hole at its tip. (2) A tungsten microwire shuttle is threaded through the guide hole and advanced into the brain. (3) As the shuttle reaches the target depth, the spiral wraps around it into a helical coil. (4) After shuttle withdrawal, the structure remains as a spring that deforms with brain growth while maintaining electrode position and contact. **b**, Exploded schematic of the multilayer device architecture consisting of two Cr/Au/Cr interconnect layers, three SU-8 passivation layers and Pt-coated electrodes. **c**, Bright-field (BF) image of the as-fabricated spiral electronics on a silicon substrate. **d**, Magnified BF image of the electrode array region. **e**, Series of BF images showing implantation of a planar spiral into a brain-mimicking gel, illustrating wrapping of the flexible spiral device around the tungsten shuttle and the stable helical spring structure after shuttle withdraw. **f,g**, FEA maps of von Mises stress (MPa) in the spring interconnect during insertion **(f)** and under longitudinal elongation following the volumetric brain growth **(g)**, confirming mechanically compliant operation. **h**, Exploded view of the neonatal headstage, including a reversible protective cover, sliding guide and custom flexible printed cable (FPC) that plugs into the recording PCB to support the chronic implantation and recording in neonatal rats. **i**, Photograph of spiral-spring electronics implanted in the neonatal rat brain. **j**, Photograph showing dam acceptance after surgery; the compact headstage allows pups to be cohoused with the dam until weaning. **k**, Photographs of representative implanted rats at key development stages from implantation at P6, first recording at P10, eye opening at P14, after weaning at P23, and young adulthood at P45.

For longitudinal recording throughout brain development, the device incorporates three key design features (Fig. 2b, Extended Data Fig. 2a): (i) flexible, spiral interconnects consisting of two Cr/Au/Cr conductive layers (3/50/3 nm) encapsulated between three SU-8 layers of 5 µm thickness; (ii) platinum (Pt)-coated high-density microelectrode arrays for simultaneous recording of single-neuron activity by multiple nearby electrodes, allowing for precise single-unit isolation and alignment^29^; and (iii) two layers of electrode arrays with a total of 32 electrodes per spiral, positioned near the inner end of the spiral structure. Fig. 2c shows a bright field (BF) image of a fabricated planar spiral device. Each device contains 32 Pt electrodes with Au interconnects (Fig. 2d). Notably, to maintain structural integrity, we incorporated twist-off support bridges between spiral turns. These temporary supports stabilize the millimeter-scale spiral during release, handling, and surgical transfer. Scanning electron microscopy (SEM) images confirm the device’s spiral structures with electrode openings, and cross-sectional SEM images show the double Au interconnect layers fully encapsulated in SU-8 passivation layers (Extended Data Fig. 2b-d).

During stereotaxic implantation, a fine tungsten shuttle wire is threaded through a guide hole at the inner end of the device. To improve mechanical robustness, the inner end of the spiral is locally reinforced with doubled Au layers wherever possible without risking electrical shorting. And the guide holes are patterned slightly smaller than the tungsten shuttle wire to ensure a snug fit during implantation. As the device advances into the tissue, the planar spiral wraps around the shuttle and self-assembles into a helical spring. After the shuttle is withdrawn, the spring configuration remains stable within the brain (Fig. 2e). FEA using the real spiral-to-spring geometry indicated that both implantation (Fig. 2f) and subsequent development-related elongation (Fig. 2g), modeled as up to 30% longitudinal elongation, produced a maximum stress of ∼40 MPa in the SU-8 ribbons, well below the reported yield strength of SU-8 (ca. 80 MPa)^30^. Next, we evaluated the electromechanical properties and stability of the device (Extended Data Fig. 3a,b) after transformation in a swollen, brain-phantom gel, capable of deforming like the developmental brain. Impedance measurements confirmed robust electrical performance: fully encapsulated interconnects maintained high conductivity after one month of implantation (Extended Data Fig. 3c). The devices also remain mechanically intact during release, implantation, and stretching in the gel model (Extended Data Fig. 3d).

Finally, we evaluated spiral-to-spring implantation in neonatal rats for chronic *in vivo* recordings throughout early postnatal development. A key practical challenge for postnatal implantation is that pups must be returned to the dam for maternal care to ensure normal development and high survival, yet the dam often chews exposed I/O connectors and the headstage. To address this, we developed a neonatal surgery and housing protocol compatible with maternal cohousing, including a protective, reversible headstage cover that preserves headstage integrity (Methods). The cover uses a sliding mechanism: when closed, it shields the headstage from the dam before weaning; when slid open, it enables connection to the recording printed circuit board (PCB) (Fig. 2h). The headstage base was 3D-printed in plastic to minimize weight, while the top slider was fabricated from aluminum to resist biting while remaining lightweight. The complete assembly weighed <2 g, allowing pups to lift their heads and move freely immediately after surgery.

With the custom-designed headstage, spiral-to-spring devices were successfully implanted in P6 rat pups. Figure 2i shows a representative implantation, where two devices were implanted in the right hemisphere, targeting V1 and mPFC, after which the I/O connects were secured under the custom headstage. Importantly, pups implanted with the device and fitted with the protective cover were accepted and nurtured by the dam (Fig. 2j). The headstage remained stably fixed throughout development, including after weaning (P21) and into adulthood (P45) (Fig. 2k).

### Same-neuron mapping during brain development

All these design elements establish a stable platform for chronic implantation and longitudinal electrophysiological recordings across neonatal brain development. We began recording at P10.

Animals were awake, head-fixed using the custom headstage while freely running on a spherical treadmill. We simultaneously recorded local field potentials (LFPs) and single-unit activity (Extended Data Fig. 3e). The high-density electrode arrays captured activity from multiple neurons simultaneously, with spikes from each neuron detected on several neighboring channels. Using these multi-channel waveforms, we can estimate each unit’s center-of-mass location from its spatial amplitude profile (Fig. 3a, left) and compute its mean waveform following a previously established pipeline (Fig. 3a, right)^31–34^. Figure 3b,c show representative traces and spikes from one recording session.

**Fig. 3.**
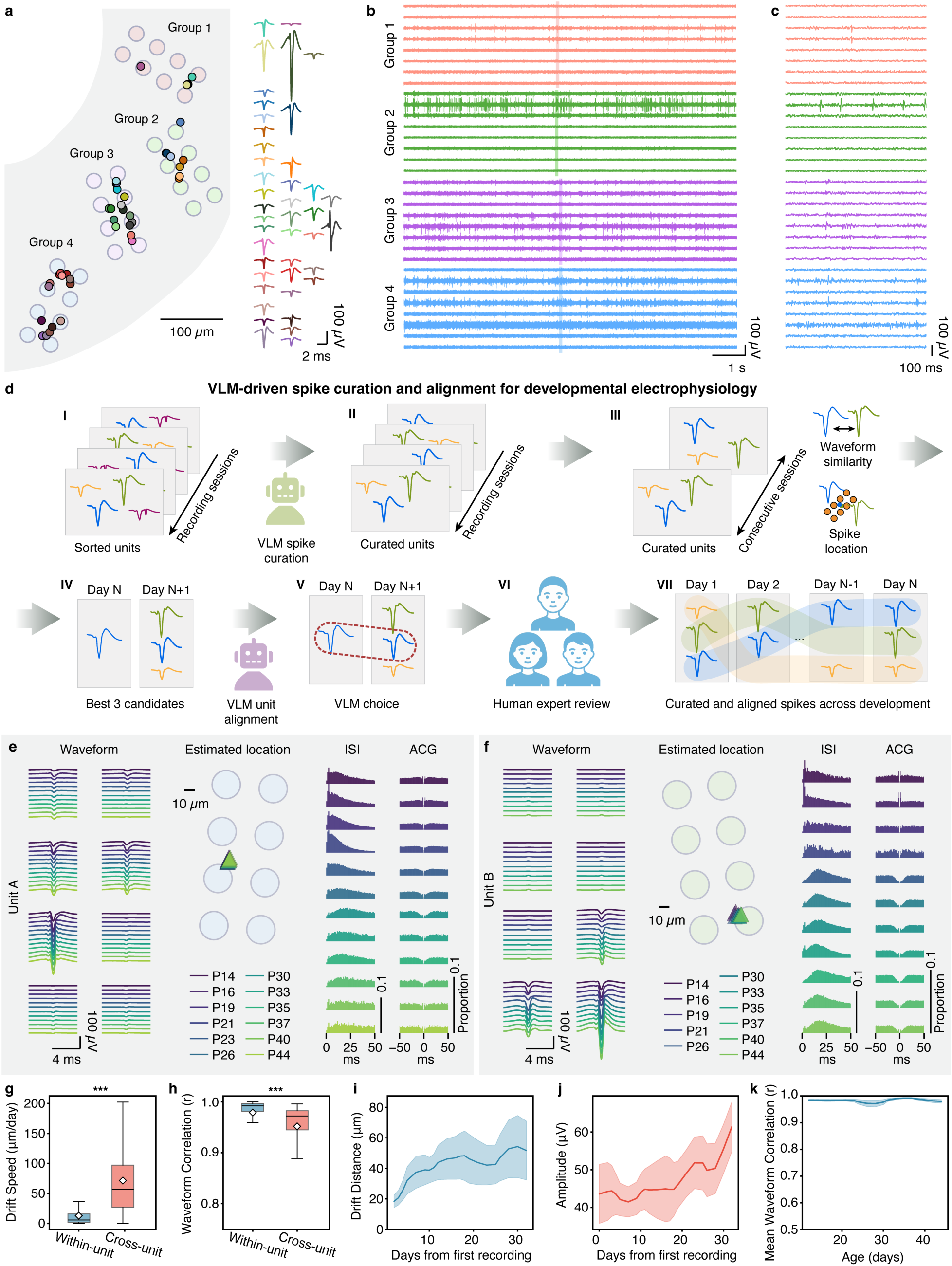
Computational pipeline for stable mapping of activity from the same neurons during brain development. **a**, Sorted single-unit waveforms (right) from representative spring electronics with four electrode arrays. Each array includes eight electrodes. The inferred location of neurons from their multi-channel waveform magnitudes across the eight electrodes are overlayed with the layout of the electrode array (left). Scale bar, 100 µm. **b,c**, Representative raw voltage traces from 32 channels (same color denotes electrodes within the same group) and a zoom-in panel **(c)** of the highlighted time shown in **(b). d**, Schematics showing Vision-language model (VLM)-driven agentic AI-human collaboration paradigm to determine and align the units across different developmental days. Spikes from each session are independently sorted and automatically curated by using a VLM assistant. Units are then aligned across sessions by unit pair-wise waveform similarity and location distance, where location is estimated via relative spike amplitudes across high-density electrode array configuration. VLM is used to unbiasedly select the best pair with three human expert reviews the choice to finalize the alignment result. **e,f**, Two representative aligned neurons across different developmental days. Their corresponding unit waveforms distribution across different channels of the electrode array (colors indicate postnatal day, P14-P44), estimated current-source locations (circles represent electrode locations, triangles represent estimated locations), inter-spike interval (ISI) histograms, and autocorrelograms (ACGs). **g**, The difference in the inferred location between units over day (drift speed) comparison between within-unit and cross-unit comparisons (***: two-sample independent t-test, p<0.001). **h**, Waveform Pearson correlation between recording sessions for within-unit (blue) versus cross-unit (red) comparisons (***: two-sample independent t-test, p<0.001). **i**, Statistical summary of cumulative drift distance (μm) as a function of days from the first recording. **j**, Statistical summary of spike amplitude (μV) shown as a function of days from first recording. **k**, Mean waveform correlation (r) across all tracked neurons as a function of postnatal age. Shaded regions represent 95% confidence intervals.

Next, we asked whether the helical spring devices could accommodate brain growth, mapping the activity from the same neurons across weeks. Conventional longitudinal alignment of single-unit signals typically relies on stability of unit waveform shape across sessions. However, in the developing brain this assumption may not hold because as neurons develop and mature, unit waveform shape and noise characteristics can also change over time. To address this, we performed spike sorting and data curation independently for each session, followed by cross-day unit alignment (see Methods).

Importantly, high-density arrays enable identification of the same unit using each unit’s spatial footprint, the characteristic pattern of spike waveforms across neighboring channels, arising from the neuron’s position relative to the array^35^. If a neuron maintains the same relative position with respect to the electrodes, its inter-channel amplitude pattern, i.e. the relative distribution of amplitudes across channels, should remain stable even when absolute amplitudes change^31^. We therefore used spatial footprint and waveform features to align single units across days.

We performed longitudinal recordings in five animals with recordings spanning development through adulthood. We consistently obtained >50 well-isolated units per animal across >10 sessions (2-3-day intervals, P10-P45). To enable unbiased, standardized, and reproducible curation, we combined an agentic AI-driven spike-sorting pipeline based on multimodal vision-language models (VLMs) and human expert review. This AI-driven pipeline, built on our recent work^32,36^, standardizes unit curation and cross-day alignment. Following that, all the results were independently verified by three human experts to ensure rigorous single-unit isolation (Fig. 3d, Methods).

For unit curation, the VLM agent evaluates units’ waveform, spatial footprint from multiple channels, estimated unit locations, inter-spike intervals (ISIs), and autocorrelograms (ACGs) to classify unit clusters as neural signals versus noise (Extended Data Fig. 4a). For cross-day probabilistic alignment, a pre-screening step was applied to identify top candidate matches based on waveform similarity and spatial proximity (estimated unit location change is <200 µm). The VLM alignment agent then assesses candidate pairs using template consistency, spatial location, and firing statistics, with final matches determined by human expert consensus (Extended Data Fig. 4b). Using this pipeline, we longitudinally mapped well-isolated units across development.

UMAP embeddings showed that multi-channel waveforms of aligned units remained stable across development, supporting successful probabilistic mapping of activity from the same neurons (Extended Data Fig. 4c)^37^.

Figure 3e-f show two representative aligned units tracked across sessions, with stable waveform shapes and spatial footprints across postnatal days, along with their corresponding spike-timing summaries (ISI and ACG). Across all units, we quantified alignment quality using three complementary readouts. First, we quantified spatial drift speed, defined as the day-to-day change in the unit’s estimated location. Within-unit spatial drift (13.0 ± 19.4 μm/day, n=896) was significantly lower than cross-unit drift (71.5 ± 60.7 μm/day, n=7,832; two-tailed t-test, p<0.001), indicating that aligned units remain spatially localized over time (Fig. 3g). Second, we quantified waveform similarity as Pearson correlation between average spike waveforms across sessions (higher values mean more similar waveforms). Within-unit correlations (0.979 ± 0.045, n=2,734) were significantly higher than cross-unit correlations (0.952 ± 0.065, n=5,367; two-tailed t-test, p<0.001), supporting the conclusion that the matched units have higher waveform similarity compared to the others across recording sessions (Fig. 3h). Third, we quantified spatial drift distance (cumulative displacement from the first recording), spike amplitude (measured spike size in µV), and session-to-session average waveform correlation, which remained stable throughout development (Fig. 3i-k), supporting reliable longitudinal single-unit mapping.

Being able to follow individual neurons across development allowed us to quantify how the firing dynamics of the aligned neurons evolve over neonatal development. Across representative units (Extended Data Fig. 5a-b) and the population (Extended Data Fig. 5c-i), we observed developmental changes consistent with prior reports of reduced bursting and strengthened post-spike suppression^38–40^. We quantified six spike-train metrics, whose age-dependent trends collectively support a transition from globally synchronized activity toward more decorrelated firing and more locally structured network dynamics^4,14,39^. Five metrics showed significant monotonic decreases with age (Spearman rank correlation, two-tailed tests): (1) Burst index, which captures the tendency for spikes to cluster into brief bursts (higher values indicate more bursty firing), decreased with age (Extended Data Fig. 5c; ρ=-0.12, p=1.62×10^-2^, n=396); (2) Coefficient of variation (CV2), a local measure of irregularity computed from successive ISIs (lower values indicate more regular firing), decreased with age (Extended Data Fig. 5e; ρ=-0.28, p=1.15×10^−8^, n=396); (3) Local variation (Lv), another ISI-based regularity metric (lower values indicate more regular firing), decreased with age (Extended Data Fig. 5f; ρ= -0.29, p=4.30×10^-9^, n=396); (4) Fano factor, defined as variance-to-mean ratio of spike counts across time windows (higher values indicate greater spike count variability), decreased with age (Extended Data Fig. 5g; ρ=-0.34, p=1.83×10^-12^, n=396); (5) Mean pairwise correlation, defined as the average Pearson correlation of 50 ms-binned spike counts across all simultaneously recorded neuron pairs within a session (higher values indicate stronger shared fluctuations), decreased with age (Extended Data Fig. 5h; ρ=-0.36, p=1.58×10^-13^, n=396).

In addition, ACG trough depth, which reflects the post-spike dip in the ACG (more negative values indicate stronger post-spike suppression), showed a significant upward trend toward zero (Extended Data Fig. 5a-b<u>,d</u>; ρ=0.33, p=4.08×10^-12^, n=418). Consistent with these statistics, session-wise spike correlation matrices evolved from broadly positive, structured correlations at early ages to weaker and more heterogeneous patterns at later ages (Extended Data Fig. 5i), indicating progressive decorrelation during cortical maturation.

### Same-neuron mapping reveals bimodal coupling states and trajectory-specific maturation

A unique advantage of our approach is its ability to follow the same neurons across weeks of rapid postnatal brain growth. Most prior studies of early postnatal circuit maturation rely on cross-sectional sampling, which records different neurons at different ages and often across different animals, and then infer developmental programs from population statistics. Without the ability to track activity from the same neurons, these approaches cannot determine whether observed changes reflect uniform changes across all neurons or within specific neuronal subpopulations. By enabling chronic, stable, single-neuron-resolved recordings from P10 to P45, our platform cannot only address this question, but also open previously inaccessible questions, such as how individual neurons contribute to population decorrelation during development. Do all neurons gradually reduce correlations in a similar manner, or do distinct subpopulations follow different developmental trajectories?

To address this, we quantified how each neuron coordinated with population-wide activity using population coupling, defined as the correlation between a neuron’s firing rate and the mean activity of the surrounding population (Fig. 4a). In adult cortex, previous cross-sectional studies have reported coexisting “soloist (low population coupling)” and “chorister (high population coupling)” neurons^41–43^. However, whether these two states are fixed cellular properties established early in development, or emerge through developmental transitions, has remained unknown (Fig. 4b). Leveraging the ability of helical spring electronics to track activity from the same neurons throughout development, we directly tested these competing hypotheses: uniform population coupling reduction across all neurons versus neuronal subgroup-specific changes (Fig. 4c)?

**Fig. 4.**
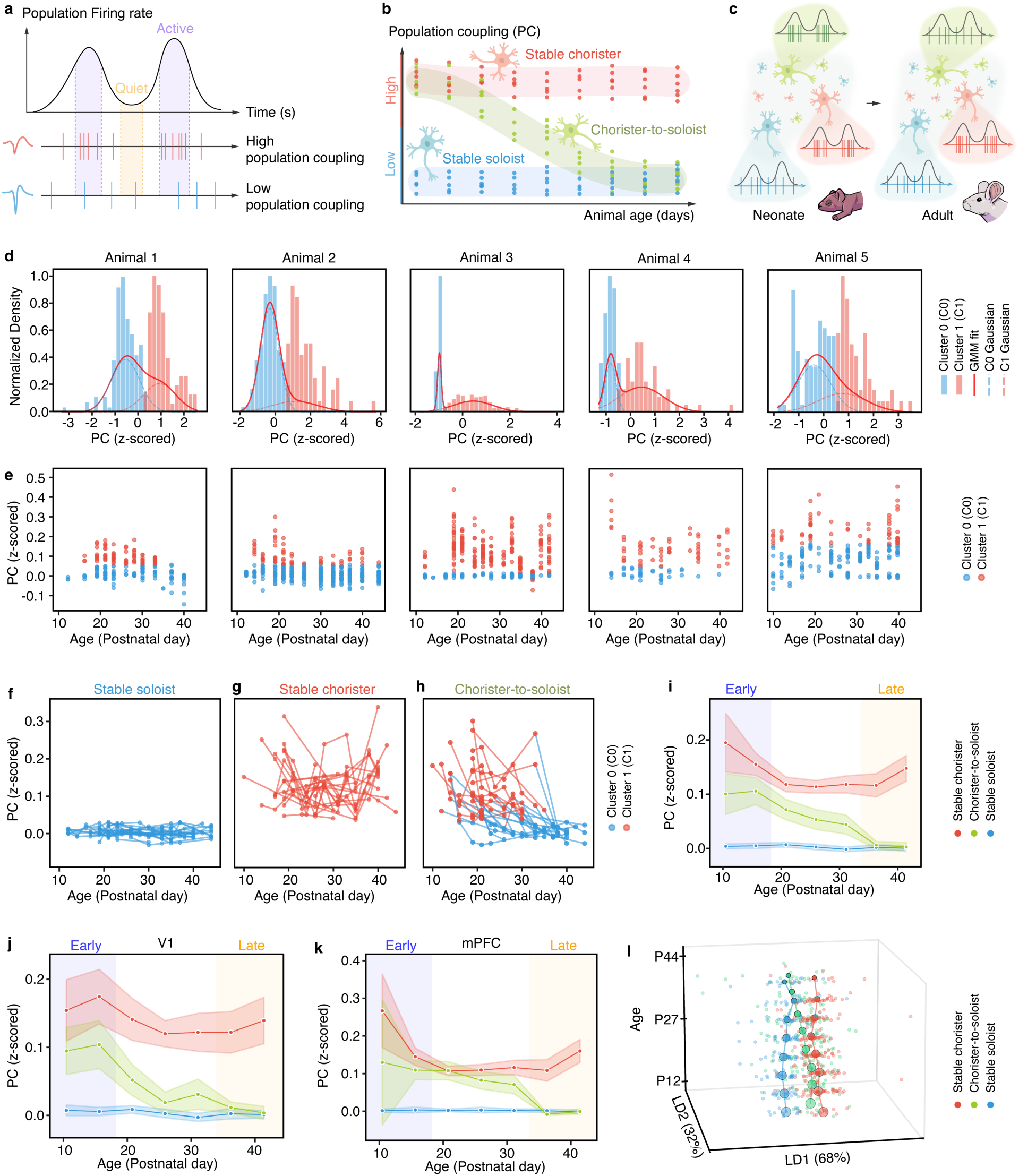
Population coupling reveals trajectory-specific network coordination strategies. **a**, Schematic illustrating population coupling (PC) measurement. Top: population firing rate fluctuations with periods of elevated activity (purple shading) and periods of lower activity (yellow shading). Bottom: example neurons with high PC (red) and low PC (blue) relative to population activity are shown below. **b**, Conceptual diagram showing trajectory-specific PC patterns across development. **c**, Schematic explaining the chorister-to-soloist transition occurring in neonatal brain. **d**, PC distributions for five animals showing bimodal structure. Blue bars: Cluster 0 (low PC). Red bars: Cluster 1 (high PC). Dashed lines: individual Gaussian components. Solid red line: Gaussian Mixture Model (GMM) fit. **e**, Individual neuron PC values across development for five animals. Each dot represents one observation. Blue: Cluster 0 membership; red: Cluster 1 membership**. f-h**, Single-neuron PC trajectories by developmental type. **f**, Stable soloists maintain stable low PC. **g**, Stable choristers maintain stable high PC. **h**, Chorister-to-soloists show systematic reduction from high to low PC. Individual trajectories are shown as thin lines. **i**, PC trajectories for all three types across entire dataset. Lines show mean values; shaded regions indicate error bands. Background shading indicates early and late developmental periods. **j,k**, Region-specific PC trajectories. **j**, Visual cortex. **k**, Medial prefrontal cortex. Lines show mean values; shaded regions indicate error bands. Background shading indicates developmental periods. **l**, Linear discriminant analysis (LDA) visualization of neuronal functional states throughout developmental timepoints. Each age point shows neurons from all three trajectory types embedded in the same 2D LDA (LD1-LD2) space computed across all timepoints. Individual neurons are shown as dots and trajectory centroids are marked with semi-transparent circles for each age bin.

We first verified whether two coupling states coexist across all ages based on stable recording data. To do so, we pooled population-coupling values across all recording days for each animal. Strikingly, coupling histograms were consistently bimodal across all animals (Fig. 4d), suggesting two functional participation states. To quantify this structure, we fit Gaussian mixture modeling (GMM), which provides a probabilistic framework for identifying overlapping subpopulations when the underlying distribution is a mixture of approximately Gaussian components^44,45^. Model selection using Bayesian Information Criterion (BIC), which penalizes additional components to favor the most parsimonious explanation^46^, together with cross-validation, strongly supported k=2 as optimal (Extended Data Fig. 6a). Across k=2-5, the two-component model achieved the lowest BIC, indicating the best trade-off between goodness-of-fit and model complexity. Five-fold cross-validation using logistic regression achieved high classification performance (CV AUROC=0.946), silhouette analysis showed clear cluster separation (silhouette coefficient=0.570), and all animals showed separation exceeding 1σ, indicating well-resolved cluster centroids (Extended Data Fig. 6b-d). Together, these metrics consistently supported two distinct populations: low-population coupling (Cluster 0) and high-population coupling (Cluster 1), reproducibly identified across all animals.

We next evaluated whether the Gaussian assumption underlying GMM was reasonable within each cluster. Shapiro-Wilk tests, quantile-quantile (Q-Q) plots, and residual analyses indicated that both cluster distributions were well-approximated by normal distributions (Extended Data Fig. 7)^47–49^. Q-Q plots showed a near-linear fit (R²>0.95), and residuals were approximately normally distributed, supporting the use of GMM. Bootstrap resampling (100 iterations) further confirmed exceptional cluster stability (98.0±1.8% agreement), validating the robustness of the two-state decomposition. Critically, both coupling states coexisted at every developmental age: neither state emerged nor disappeared during development (Fig. 4e). This demonstrates that coupling heterogeneity is not a product of maturation but rather a fundamental organizational principle present from early circuit formation.

### Same-neuron mapping reveals three developmental trajectories

While the bimodal population states persisted over development, following the history of each individual neuron revealed heterogeneity in how each neuron’s coupling changes over time. Based on each neuron’s day-by-day cluster assignment, we identified three distinct developmental trajectories. The first trajectory was what we call stable soloist neurons, which maintained consistently low population coupling (Cluster 0) throughout development (Fig. 4f). The second trajectory was stable chorister neurons, which maintained consistently high coupling (Cluster 1) throughout development (Fig. 4g). Most importantly, we identified chorister-to-soloist neurons, a third trajectory, which showed a systematic transition from high population coupling (Cluster 1) to low population coupling (Cluster 0) throughout development (Fig. 4h). These groups and trajectories cannot be identified in cross-sectional data.

Averaging population coupling within each trajectory confirmed distinct developmental patterns (Fig. 4i). Stable soloist neurons maintained near-zero population coupling (0.00-0.01) across P10-P45 with no significant trend (Mann-Kendall p=0.133, Spearman ρ=-0.750, p=0.052). Stable chorister neurons remained elevated and stable (0.11-0.20) across P10-P45 with also no significant trend (Mann-Kendall p=0.368, Spearman ρ=-0.464, p=0.294). In contrast to the other two trajectories, chorister-to-soloist neurons showed progressive decoupling, significantly declining from intermediate-high coupling (0.10 at P10-P14) to stable soloist-like levels (0.00 at P35-P40), representing an approximately 95% reduction in synchronization with population activity (Mann-Kendall p=0.007, Spearman ρ=-0.964, p=0.0005). While all three groups were recorded across most ages (Extended Data Fig. 8a), the chorister-to-soloist transition predominantly occurred during P21-P35, coinciding with established critical periods for cortical refinement (Extended Data Fig. 8b) ^50–53^.

These tripartite development trajectories are accessible only through long-term same-neuron mapping at single-cell resolution. Conventional cross-sectional studies collect data from different animals at each age, treating each measurement as independent. This approach cannot distinguish neurons that actively transition between coordination states (chorister-to-soloists) from neurons that maintain stable coupling throughout development (stable soloists and stable choristers).

Cross-sectional analysis can only report the aggregate distribution of population coupling values at each age (Extended Data Fig. 8c), which shows an apparent uniform developmental decline (r=-0.174, R²=3.0%, p<0.001, n=741 observations across P10-P45). However, continuous longitudinal probabilistic mapping of the same neurons reveals that this aggregate pattern conceals three distinct developmental trajectories. Variance decomposition analysis demonstrates that the cross-sectional view fundamentally misrepresents the underlying biology (Extended Data Fig. 8d). Trajectory identity alone explains 45.3% of population coupling variance (R^2^=0.453, η²=0.453, F(2,738)=306.1, p=1.6×10⁻⁹⁷), a 15.2-fold increase over age alone (R^2^=0.030). The three trajectories are separated by large effect sizes (Cohen’s d: Stable soloist vs. stable chorister = -2.36, stable soloist vs. chorister-to-soloist = -1.01, stable chorister vs. chorister-to-soloist = 1.14), indicating they represent categorically distinct neuronal populations rather than continuous variation (Extended Data Fig. 8e). The additive model including age and trajectory identity (R^2^=0.484, AIC = -2151.6) substantially outperforms age-only models, confirming that developmental trajectories, not chronological age, are the primary determinant of population coupling.

We next asked whether these trajectories differ across cortical regions. When pooling population coupling values across ages and separating units from V1 and mPFC, mPFC showed a larger fraction of high population coupling neurons (Extended Data Fig. 9a-c). Trajectory composition also differed, showing that mPFC was dominated by stable choristers (58%), whereas V1 did not (Extended Data Fig. 9d,e). Moreover, the timing of chorister-to-soloist transitions was region-dependent. In V1, decoupling occurred primarily during P15–P27, whereas in mPFC transitions were delayed to P28–P45 (Fig. 4j,k). This 10-day offset aligns with known developmental gradients in which sensory cortices refine earlier than association cortices^54–57^.

### Independent metrics confirm transformation across organizational levels

The discovery that chorister-to-soloist neurons transition between network coordination states raised a fundamental question: is this transition specific to population coupling, or does it reflect a broader maturation of neuronal function? To address this, we compared electrophysiological properties across the three trajectory groups using metrics that are independent of population coupling. For example, firing irregularity, quantified by Lv of ISIs decreased significantly in stable choristers (-10% from 1.18 to 1.06, Mann-Kendall p=0.003, Spearman ρ=-1.00, p<0.0001) and chorister-to-soloists (-19%, from 1.15 to 0.92, Mann-Kendall p=0.036, Spearman ρ=-0.857, p=0.014), but remained stable in stable soloists (-12%, Mann-Kendall p=0.133).

Beyond the Lv of ISIs, the three population coupling trajectories diverged across a broader set of electrophysiological properties. We quantified 14 electrophysiological metrics spanning spike waveform, intrinsic firing statistics, spike train variability, oscillatory coupling strength, and autocorrelation structure, none directly related to population coupling (Methods; Extended Data Fig.<u>10a</u>). Although trajectory membership was defined solely by population coupling dynamics, each group exhibited distinct metric profiles and developmental trends.

To summarize these multivariate differences, we projected neurons into a 2D linear discriminant analysis (LDA) space (LD1-LD2) computed from all 14 metrics^58,59^. (Extended Data Fig.<u>10b,c</u>).

Separation in this space was dominated by waveform and excitability-related features: LD1 captured 68.4% of between-group discriminative variance and was driven most strongly by repolarization slope (+1.81), peak-to-peak amplitude (-1.14), firing rate (-0.98), Fano factor (-0.63) (Extended Data Fig.<u>10d</u>). LD2 captured the remaining 31.6 % and was driven by peak-to-peak amplitude (-1.28), repolarization slope (+0.93), firing rate (-0.84), and gamma phase locking strength (-0.43) (Extended Data Fig.<u>10e</u>). Together, LD1 and LD2 captures 100% of the discriminative variance separating the three trajectories.

Consistent with the metric profiles, stable soloists and stable choristers occupied distinct regions of LD space across development. Repolarization slope was higher in stable soloists than in stable choristers across age (Extended Data Fig. 10a), aligning with the negative loading of peak-to-peak voltage on LD1. Several waveform- and firing-related measures also showed large between-group effect sizes between stable soloists and stable choristers, indicating robust separation of these groups in distinct biophysical/excitability regimes rather than just in population coupling values.

Across development, the three trajectories also differed in which metrics changed most strongly with age. For stable soloists, waveform asymmetry increased with age (r=0.405) and peak-to-peak voltage increased modestly (r=0.157), while most other metrics remained comparatively stable. For stable choristers, repolarization slope increased (r=0.254), and peak-to-peak voltage increased (r=0.176), alongside decrease in Lv (r=-0.213). Chorister-to-soloists showed the largest set of age-dependent changes, including strong decreases in Fano factor (r=-0.464), trough-to-peak width (-0.298), Lv (r=-0.286), and burst index (r=-0.255). The 2D LDA space (LD1-LD2) projection visualizes these group-specific developmental trends, showing relatively stable centroid positions for stable soloists and stable choristers across development, contrasting with systematic chorister-to-soloist migration (Fig. 4l). Chorister-to-soloists began in stable chorister-proximal functional space at P10, traversed intermediate configurations during P21-P31, and converged toward stable soloist-proximal space by P40-P45. The smooth, continuous path indicates gradual functional reorganization rather than discrete state-switching.

## Discussion

A central challenge in developmental electrophysiology is how to maintain a stable, single-neuron interface while the brain undergoes rapid growth, displacement, and deformation. This growth-adaptive brain-machine interface presented here addresses this limitation by coupling standard planar microfabricated flexible electronics with an *in situ* spiral-to-spring transformation during implantation. The resulting helical spring architecture provides low axial stiffness and high compliance along the direction of constraint imposed by skull anchoring, while preserving stable local contact between tissue and recording sites. Together with an analysis workflow tailored to the developing brain, combining session-wise spike sorting with cross-session probabilistic cross-session alignment using high-density spatial waveform footprints, this platform enables continuous, spike-resolved recordings from the same neurons across weeks of early postnatal development *in vivo*.

With this capability, we could move beyond population-averaged characterization of cortical maturation to directly test how correlations between individual neuron firing and population activity change across development. Early cortical networks are known to exhibit large, highly synchronous activity events that gradually give way to sparser, more decorrelated firing patterns as circuits mature^3,4^. Prior work has implicated inhibitory maturation and shifts in excitation-inhibition balance as sufficient drivers of this transition at the population level^60^. However, because prior studies rely on cross-sectional sampling not from the same neurons, or at best relatively short longitudinal windows, they cannot determine whether developmental desynchronization reflects a broadly shared adjustment across most neurons or a selective reorganization in which only specific subpopulations change their role while others remain stable^14,60^.

Here, long-term same-neuron mapping throughout development reveals that developmental desynchronization in rat visual cortex and mPFC is driven primarily through trajectory-specific reorganization of a subset of neurons, rather than uniform changes across the population. Using population coupling as a compact measure of each neuron’s coordination with local population activity, and building on the “chorister/soloist” framework established in adult sensory cortex^43^, we identify two stable participation states that persist at all ages: neurons that remain weakly coupled throughout development (stable soloists) and neurons that remain strongly coupled throughout development (stable choristers). Critically, the same-neuron mapping uncovers a third state that is invisible to cross-sectional studies: neurons that begin in a chorister-like state but progressively reduce their population coupling over the third to fifth postnatal weeks, becoming soloist-like by adulthood (chorister-to-soloists). At the network level, the developmental decline in mean population coupling and pairwise correlations from P10 to P45 is largely explained by this transitioning subgroup, whereas stable soloists and stable choristers show comparatively minor changes. Thus, the well-described shift from synchronous to decorrelated activity emerges, in these circuits, because a defined subset of neurons changes its relationship to the local population, while other neurons preserve their participation state.

Interestingly, the trajectories defined purely by population coupling dynamics are not confined to coupling-related measures, but are accompanied by coordinated changes in electrophysiological properties that extend beyond coupling itself. The chorister-to-soloist group shows the largest age-dependent changes across multiple metric classes, including spike-train variability, burst statistics, oscillatory coupling strength, and spike waveform features. This pattern aligns with prior studies linking developmental decorrelation and changes in burst structure, rhythmic coupling, and spike-train regularity^14,60^. By contrast, the two stable trajectories exhibit more limited or gradual changes across many metrics, indicating that developmental reorganization is not uniform across all neurons.

These observations intersect naturally with critical-period concepts and provide a structured way to examine developmental timing. The interval right after eye opening is a canonical time window of rapid circuit reorganization^53,61,62^, and in our data the chorister-to-soloist group shows its strongest coordinated changes during this interval. Prior theoretical and computational works in adult cortex propose highly coupled neurons may constitute a more plastic, population-locked substrate while weakly coupled neurons support more stable, individual-specific representations^43,63,64^. Consistent with this, our results further support a developmental picture in which only a subset of initially strongly coupled neurons is progressively reassigned toward a more independent operating mode as circuits mature. Chorister-to-soloist neurons exhibit the strongest reduction in burst firing, a transient elevation in coupling to gamma-frequency activity during the same transition window, and pronounced maturation of action potential waveform kinetics. Notably, the trajectories do not simply reflect uniform maturation across all neurons: stable soloists and stable choristers remain distinguishable across development in the multivariate physiological space, while the transitioning group shows the strongest movement.

These findings also refine how critical periods and neurodevelopmental disorders may be evaluated. Many models are commonly summarized using population-averaged descriptors such as hypersynchrony, altered excitation-inhibition balance, or reduced decorrelation. However, such readouts cannot distinguish global shifts from selective disruption of specific developmental patterns. In genetic or environmental perturbation models relevant to autism spectrum disorder or schizophrenia, the trajectory approach suggests a concrete set of tests as next steps: whether one or more trajectories are reduced or overrepresented, whether the chorister-to-soloist transition timing is delayed or altered, and whether the accompanying maturation of firing, variability, and waveform metrics are preserved. Integrating trajectory-based analyses with assays of inhibitory maturation and circuit state could help determine whether observed phenotypes reflect broad changes affecting all neurons versus changes concentrated within specific developmental trajectories as causal pathways.

More broadly, growth-adaptive, depth-wise stretchable electronics offer a general strategy for stable electrophysiological recording in dynamically deforming tissues. Similar design principles may extend chronic, high-resolution recordings to other developing organs or mechanically active systems such as the heart. In the developing brain, this platform shifts the unit of description from population-level decorrelation to identifiable single-neuron trajectories that implement it. Combining growth-adaptive recordings with molecular tagging, cell-type-specific recording and perturbations, and broader deployment across regions and species should clarify the generality of trajectory-structured maturation and provide a tractable target for understanding, and ultimately correcting, disrupted development.

## Data availability

The authors declare that data supporting the findings of this study are available from the corresponding author upon request.

## Acknowledgments

J.L. acknowledges the support from NIH/NIMH 1RF1MH123948; NIH/NICHD 1R01HD115272; NSF/EFRI 2422348, National Research Foundation of Korea grant funded by the Korea government (MSIT) (RS-2024-00460364), DGIST R&D Program of the Korean Ministry of Science and ICT (25-IRJoint-01), and NSF through the Harvard University Material Research Science and Engineering Center (DMR-2011754). A.L. acknowledges that this material is based upon work supported by the NSF Graduate Research Fellowship Program under Grant No. (1745302 and 2141064). H.S. acknowledges the support from Aramont Fund for Emerging

Science Research. A.M.L. acknowledges the support from the RCC-Fellowship of Harvard University and the Excellence Fellowship of the Fundacion Rafael del Pino.

## Author contributions

H.S., A.L., and J.L. conceived the idea. A.L. and H.S. fabricated and characterized electronics. A.L., H.S., and J.Lee performed device characterization. A.L. and H.S. performed implantation and electrical recording. Z.W. did the mechanical simulation. A.L., H.S., A.M.L., and X.Z. analyzed data. A.L., H.S. and A.M.L. prepared figures and wrote the manuscript. All authors reviewed the figures and the manuscript. J.L. supervised the study.

## Additional information

Correspondence and requests for materials should be addressed to Jia Liu.

## Methods

### 1. Chemicals and reagents

All chemicals were obtained from Sigma-Aldrich unless otherwise mentioned and used without further purification. All photoresists and developers in the nanofabrication were obtained from MicroChem Corporation unless further notification.

### 2. Fabrication of spring electronics

#### Cleanroom fabrication

(1) Wafer cleaning: A 3-inch thermal oxide silicon wafer (2005, University wafer) was rinsed with acetone, isopropyl alcohol (IPA), water, then blown dry and baked at 110 °C for 3 minutes, followed by O_2_ plasma treatment at 100 W, 40 sccm O_2_ for 5 minutes. (2) Ni sacrificial layer: Hexamethyldisilazane (HMDS) was spin-coated at 3,000 rpm for 1 minute. LOR 3A was spin-coated at 3,000 rpm for 1 minute and hard-baked at 180 °C for 5 minutes. S1805 was spin-coated at 3,000 rpm for 1 minute and hard-baked at 115 °C for 1 minute. Then the photoresists were exposed with 40 mJ/cm^2^ ultraviolet (UV) light and developed with CD-26 for 1 minute, rinsed with deionized (DI) water, and blown dry. After preparing the photoresist pattern, a 100 nm Ni layer was thermally deposited on the wafer (Sharon) and lifted off in Remover PG for 3 hours. (3) Bottom SU-8 passivation layer: SU-8 2002 was spin-coated twice, at 2,200 rpm for 1 minute and pre-baked at 60 °C for 1 minute and 95 °C for 4 minutes. SU-8 was exposed with 200 mJ/cm^2^ UV light, then post-baked at 60 °C for 1 minute and 95 °C for 2 minutes. SU-8 was then developed in an SU-8 developer for 1 minute, rinsed with IPA, and blown dry. Finally, SU-8 was hard-baked at 180 °C for 30 minutes. (5) Bottom Au interconnects: 3/50/3 nm Cr/Au/Cr layers were deposited on the top of the bottom SU-8 passivation layer by e-beam evaporator (Denton) with S1805 and lifted off in Remover PG for 8 hours. (6) Middle SU-8 passivation layer: The fabrication of the middle SU-8 layer followed the same procedure as the fabrication of the bottom SU-8 layer. (7) Top Au interconnects: The fabrication of the top Au interconnects layer followed the same procedure as the fabrication of the bottom Au layer. (8) Pt layer: 3/40 nm chromium (Cr)/Pt layers were deposited by electron-beam (e-beam) evaporator (Denton) with photoresist and lifted off in Remover PG for 2 hours. (9) Top SU-8 passivation layer: The fabrication of the top SU-8 layer followed the same procedure as the fabrication of the bottom and middle SU-8 layer.

#### Device treatment for implantation

(1) Wafer was cut using a dicing saw, protected by photoresist S1813 during the cutting process, and the photoresist was removed afterwards. (2) A flexible flat cable (FFC) (Molex) was bonded to the I/O pads using a flip-chip bonder (Finetech Fineplacer). (3) The device was released from the substrate by removing the Ni sacrificial layer in Ni etchant (TFB, Transene Company), which was followed by washing with DI water. The device was then disinfected by incubating with IPA solution overnight.

### 3. Characterizations of spring electronics

#### Mechanical simulation

All FEA were performed using ABAQUS 2024 and 2025. Following material properties are used: *E* = 6 GPa and *v* = 0.22 for SU-8, *E* = 0.5 MPa and *v* = 0.49 for PFPE-DMA, *E* = 79 GPa and *v* = 0.42 for Au, *E* = 0.5 MPa and *v* = 0.49 for tissue. Concentration of Cr was neglected. The ribbon was modeled as an anisotropic composite material with the effective elastic properties calculated according to the modulus and thickness of each encapsulating layer and conductive layers.

(1) Development simulation. PFPE-DMA and SU-8 ribbons were discretized using S4R shell elements while the tissue was discretized with C3D8R solid elements. Perfect bonding between the device and tissue was assumed by embedding the device elements within the tissue elements. Development was simulated through thermally induced expansion of the embryonic tissue, with thermal strain defined as 𝜀 = 𝛼Δ*T*, where 𝛼 is the artificial coefficient of thermal expansion and Δ*T* is the artificial temperature change relative to the initial state. The top surface of the tissue was constrained in the *Z*-direction to represent mechanical confinement from the skull. Simulations were performed using the static implicit solver.

(2) Implantation simulation: for computational efficiency, the interaction between the ribbon and the tissue was idealized as an elastic foundation. This foundation effect was implemented as an equivalent body force field via a user-defined subroutine. The body force was formulated as

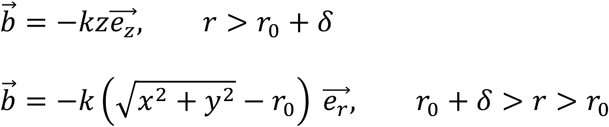

where *k* and *r*_0_are fitting parameters representing the effective stiffness of the tissue support and the equilibrium radius of the device, respectively. The parameter 𝛿 is introduced as a numerical stabilization term. The variables *x*, *y*, and *z* denote the deformed nodal coordinates of the device, while e_*z*_ and e_*r*_ are the unit vectors in the vertical (*z*) and radial (*r*) directions, respectively. A prescribed displacement in the *z*-direction was applied at the inner end of the device, with additional constraints imposed to eliminate rigid-body motion. The implantation simulation was performed using the dynamic explicit solver. (3) Stiffness simulation: to ensure force balance, a uniform traction with angle of 0°, 30°, 60°, 90° to the X axis is applied on the peripheral of the device. The directional stiffness is calculated based on the applied force on one side and the component of displacement in the direction of interest.

#### Swelling test

The test started with dissolving. 50 g acrylamide, 0.0868 N,N’-Methylenebisacrylamide, 0.0642 g Ammonium persulfate and 0.0654 g Tetramethylethylenediamine were fully dissolved in 64 g DI water as hydrogel precursor. Then the precursor was poured in a cube mode (TED PELLA, Peel-A-Way disposable histology molds, 27110) and crosslinked as a dummy in 50 °C. The devices were embedded into the dummy while crosslinking. The hydrogel cube dummy containing devices was then soaked in DI water to swell. During and after swelling, the length of the device and the volume of the hydrogel cube were measured and calculated to reflect whether the device elongated in hydrogel swelling.

#### *In vitro* impedance measurements

The released device was tested to check electrodes’ impedance before moving forward to implantation. The *in vitro* impedance measurement mimicked the *in vivo* electrophysiology condition. The device was incubated in 37 °C PBS (VWR international, 97063-660). The impedance at 1 kHz was measured using CerePlex Direct (Blackrock Neurotech) data acquisition system. Electrochemical impedance spectra (EIS) measurement of the device was conducted using SP-150 potentiostat. A sinusoidal voltage with a 100 mV peak-to-peak amplitude was applied in the frequency range of 1 Hz to 1 MHz, using a platinum counter electrode and a silver/silver chloride (Ag/AgCl) reference electrode.

### 4. Animal experiments

Neonatal rats were bred from pregnant CD rats (Charles River Laboratories INC). Rat pups were kept on a regular 12-hour light-dark cycle and housed with their mother before and after surgery and were weaned at postnatal age 21 (P21).

#### Ethical statement

All animal experimental procedures were approved by the Institutional Animal Care and Use Committee (IACUC) of Harvard under animal protocol #19-03-348.

#### Stereotactic device implantation in neonatal rats

CD rat pups underwent device implantation surgery at P6. They were placed on a customized platform for head fixation and were maintained under anesthesia with 0.5% isoflurane during surgery. The head fixation setup was adjusted to the level where bregma and lambda were at the same height and aligned laterally. For device implantation, a circular piece of skin on the skull was cut away with surgical scissors. Then, two small cranial windows were opened, and the probe was stereotaxically implanted until all electrodes were inserted to the targeted depth.

Stereotaxic coordinates for implantation were as follows, with reference to bregma. Visual cortex (V1) in P6 rats were targeted with M/L 2.5 mm, A/P -4 mm, D/V 1.02 mm, and medial prefrontal cortex (mPFC) in P6 rats were targeted with M/L 0.36 mm, A/P 1.35 mm, D/V 2.22 mm. The coordinates were determined from the postnatal rat brain atlas database^65^. A 100 µm-thick stainless-steel wire (A-M SYSTMES, 793100) was partially inserted to the different location of the brain functioning as the ground. The wafer and I/O cable of the device and the ground wire were sealed and fixed with dental cement, followed by fixation of our custom-made 3D printed headstage. After device implantation and postoperative care, the rat pups were placed back with their dam. The rat pups were administered with extended-release buprenorphine (EthiqaXR, Fidelis) on the day of surgery and Carprofen (Rimadyl, Zoetis) on the day of surgery and once per day for the following 2 days. Activity, incision area, and pain levels of each animal were monitored and recorded daily for 4 days after implantation.

#### Electrophysiology recording

All recordings were done with RHD recording system (Intan technologies). Python codes provided by Intan were used to load, view, and convert raw data files into an accessible format for data analysis. For data acquisition, Intan RHD recording headstages were connected to the implanted device through a custom-designed PCB. The recording setup was surrounded by a Faraday cage and placed on a vibration isolated table. Signals were acquired at 30 kS/s, and each recording session was conducted every Monday, Wednesday, and Friday. Recording sessions consisted of 10 minutes of spontaneous freely moving animals head fixed on a custom-made spherical treadmill. The rat pups were acclimated to the treadmill and head fixation setup 3 days before the recording. The first recording session started at P10.

### 5. Spike data processing

Electrophysiological recordings were processed offline using custom Python 3.9 code. Raw wideband traces were bandpass filtered to 300-3000 Hz and handled using SpikeInterface objects for standardization. Spike sorting for each daily session was performed with MountainSort4 via SpikeInterface, using default detection thresholds with drift correction disabled to avoid over-merging across developmentally driven waveform changes^34,66,67^. Session-wise sorting isolated session-specific noise environments.

#### Curation

Initial sorting outputs were curated using a multimodal vision-language model (VLM) assistant adapted from prior AI electrophysiology frameworks^32,36^. For each unit, the assistant received: average waveform template, multi-channel waveform snippets, estimated spike localization (center-of-mass), ISI histogram, and ACG. The VLM analysis was conducted using OpenAI’s GPT-4o with a temperature setting of 0.7. After priming with 1-3 labeled examples per category (Extended Data Fig. 3a), the VLM classified units as valid neurons or noise with textual justification. Curated units from each session advanced to alignment. All VLM decisions were verified for within-session and across-session consistency.

#### Cross-day alignment

To identify the same neuron across recording days, curated units underwent hierarchical three-stage alignment:

**(1) Basic preprocessing and candidate filtering.** For each unit, we computed: (i) spatial footprint, normalized amplitude distribution across neighboring channels, (ii) center-of-mass location, and (iii) waveform-template similarity (Pearson correlation). Units with spatial separation <200 µm and above-threshold waveform similarity were retained as candidates. For each unit, the top three matches were preserved; units without suitable candidates received no forced assignment.
**(2) VLM-assisted alignment.** A second VLM assistant received the focal unit and its 1-3 candidates. Inputs included waveform templates, multi-channel snippets, spatial coordinates, and firing-rate statistics. After priming with curated examples of true and false matches, the model issued probabilistic decisions on whether each candidate represented the same neuron. If all candidates were rejected, the neuron was recorded as absent on that day (Extended Data Fig. 3b).
**(3) Expert adjudication.** Three independent human experts reviewed all VLM decisions to ensure biological plausibility and methodological rigor. Final alignment labels were determined by consensus (majority voting). This combined approach, high-density waveforms, quantitative pre-screening, VLM inference, and expert adjudication, enabled robust probabilistic mapping of individual neurons across >10 sessions spanning P10-P45.

#### Validation of mapping stability

To validate single-neuron identity preservation across development, we computed: (i) within-unit spatial drift (change in estimated center-of-mass location per day for the same aligned neuron), (ii) cross-unit spatial drift (distance between different neurons across consecutive sessions as control), (iii) waveform Pearson correlation for within-unit versus cross-unit comparisons across all session pairs, and (iv) cumulative spatial drift, spike amplitude changes, and session-to-session correlation values for each aligned unit throughout the P10-P45 recording period. Statistical comparisons used two-tailed independent t-tests with Cohen’s d effect sizes.

### 6. GMM fitting

Gaussian Mixture Modeling (GMM) with k=2 components was performed on population coupling metric values to identify functionally distinct neuronal populations during development. Population coupling, defined as the normalized correlation between individual neurons and population activity, is a continuous measure expected to follow Gaussian distributions under the Central Limit Theorem. GMM is the maximum likelihood approach for modeling heterogeneous populations composed of Gaussian sub-populations. Data were z-score standardized within each animal to account for animal-specific baseline differences while preserving biological signals.

#### GMM model validation

Silhouette coefficients were calculated for each neuron, measuring similarity to assigned cluster versus nearest neighboring cluster (range −1 to 1). Five-fold cross-validation using logistic regression assessed prediction of cluster assignments, with an area under the ROC curve (AUROC) computed across folds. Bayesian Information Criterion (BIC) and Akaike Information Criterion (AIC) were computed for each GMM with k=2, 3, 4, and 5 components. Component separation was quantified as: separation = |µ_1_ − µ_0_| / mean (σ_0_, σ_1_).

#### Normality assessment of GMM clusters

To validate the Gaussian assumption underlying the two-component GMM, we performed normality assessments on the population coupling distributions within each cluster. Quantile-quantile (Q-Q) plots compared sample quantiles against theoretical normal distribution quantiles, with coefficient of determination (R²) computed via linear regression. Population coupling z-scores within each cluster were binned into histograms, and Gaussian probability density functions were fitted using maximum likelihood estimation of the mean (μ) and standard deviation (σ). Residuals were computed as the difference between observed values and assigned cluster’s means. Residual normality was assessed using histogram visualization, Q-Q plots, and residual-versus-fitted plots. Formal normality tests (Shapiro-Wilk, Kolmogorov-Smirnov, D’Agostino-Pearson) were applied, along with skewness and kurtosis calculations. Given large sample sizes (N > 500), we prioritized visual assessments over p-values, as large samples can reject normality for practically negligible deviations.

### 7. Trajectory classification

Neurons were classified into three developmental trajectory types based on their longitudinal cluster assignments across recording sessions: Stable soloist neurons were defined as units that remained in the low population coupling cluster (C0) across all recorded time points, maintaining consistently sparse population coupling throughout development. Stable chorister neurons were defined as units that remained in the high population coupling cluster (C1) across all recorded time points, maintaining consistently strong population coupling throughout development.

Chorister-to-soloist neurons were defined as units that transitioned from the high-population coupling cluster (C1) to the low-population coupling cluster (C0) during development, representing neurons that underwent a developmental refinement of their population coupling properties. The trajectory analysis is based on 1258 observations with 298 total units. This classification yielded 37 Stable soloists, 67 Stable choristers, and 46 Chorister-to-soloists from 298 total tracked neurons. Neurons lacking sufficient early or late data (n=134) or showing transitions from C0 to C1 (n=14, 3.3-fold rarer than chorister-to-soloist) were excluded from trajectory analyses but retained for cross-sectional cluster characterization. Since all neurons were retained for cross-sectional cluster characterization or population-level analyses, exclusion of these neurons did not alter the findings or conclusions.

#### Variance decomposition analysis

To quantify the relative contribution of age versus trajectory identity to population coupling variance, we fit ordinary least squares regression models: (1) Age-only model: value ∼ age, (2) Trajectory-only model: value ∼ C(trajectory), (3) Additive model: value ∼ age + C(trajectory), and (4) Interaction model: value ∼ age × C(trajectory). The coefficient of determination (R²) was computed for each model to assess explained variance.

Model comparison was performed using Akaike Information Criterion (AIC) and Bayesian Information Criterion (BIC), with lower values indicating better model fit after penalizing complexity.

#### Statistical analysis of trajectories

For each trajectory type, Pearson correlation coefficients were computed between postnatal age and population coupling values to assess within-trajectory developmental trends. Effect sizes were quantified using eta-squared (η²) for the overall trajectory effect from one-way ANOVA, and Cohen’s d for pairwise trajectory comparisons.

Population coupling values were binned into 5-day age windows (P10-15, P15-20, etc.) for visualization, with mean ± SEM computed within each bin.

#### Metrics analyzed

Analyses were performed using custom written code. Burst index was calculated as the proportion of spikes occurring in bursts, defined as the fraction of ISIs < 6 ms. ACGs were computed for each neuron using 1 ms bins over a ±50 ms window. ACG trough depth was measured as the peak height (0-1 ms bin) minus the minimum value in the 2-10 ms trough region. Higher trough depth indicates a stronger refractory period. CV2, the local coefficient of variation of ISIs, was calculated as 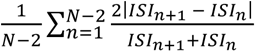 (Spike times are *t*_1_, …, *t*_*N*_. *ISI*_*n*_ = *t*_*n*+1_ − *t*_*n*_ for *n* = 1, …, *N* − 1.), i.e., averaged across consecutive ISI pairs, providing a local measure of firing irregularity that is robust to slow rate changes. Lv, the local variation of ISIs, was calculated as the mean of 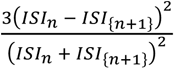 across consecutive ISI pairs.

Fano factor was computed as the variance-to-mean ratio of spike counts in 50 ms bins during spontaneous activity. For pairwise correlation analysis, spike count Pearson correlation values were computed between all simultaneously recorded neuron pairs using 50 ms bins. Correlation matrices were constructed with neurons ordered by unit number, and normalized to the range [-1, 1]. Mean pairwise correlation was calculated as the average of all unique off-diagonal elements in each session’s correlation matrix. The strength of phase-locking between spikes and the gamma band was calculated by the mean vector length (MVL) that ranges from 0 (no phase-locking) to 1 (perfect phase-locking). The spike times were converted to phase indices based on the LFP sampling rate, and the phase at each spike time was extracted from the band passed gamma LFP signal (30-80 Hz). The resultant vector’s length was defined as MVL.

#### Developmental trajectory heatmap analysis

To visualize how electrophysiological properties change across development for each trajectory type, we constructed heatmaps using 14 independent metrics that exclude population coupling-derived measures (which were used to define trajectories) to avoid circular reasoning. The 14 metrics comprised: waveform features (spike width, trough-to-peak width, repolarization slope, peak-to-peak voltage, waveform asymmetry), firing properties (firing rate, burst index, median ISI), spike timing variability (Lv, Fano factor), oscillatory phase-locking strength (gamma, beta, theta coupling), and autocorrelation feature (ACG trough depth). Data were grouped into 2-day age bins spanning P10-P44. For each trajectory type (stable soloist, stable chorister, chorister-to-soloist) and age bin, we computed the mean metric value across all observations. To reduce noise while preserving developmental trends, we applied Gaussian smoothing (σ=1.5) along the age axis, with linear interpolation to handle missing values prior to smoothing. Each metric (row) was then z-score normalized across all trajectory-age combinations (i.e., across the entire row spanning all three trajectory panels), using the formula: Z = (x − μ) / σ, where μ and σ are the mean and standard deviation of all valid values in that row.

#### Linear discriminant analysis

We used linear discriminant analysis (LDA)^58,59^ to quantify separation among the three pre-defined trajectories (stable soloist, stable chorister, and chorister-to-soloist) using 14 metrics: spike waveform width, trough-to-peak width, repolarization slop, peak-to-peak voltage, waveform asymmetry, firing rate, burst index, median ISI, Lv, Fano factor, gamma, beta, and theta coupling strength, and ACG trough depth. Data were analyzed across eight developmental windows spanning ages 10-45 days. For neurons with multiple sessions within a window, metric values were averaged per neuron within that window. Neurons were retained if they had valid values for ≥70% of metrics (≥9/14), and remaining missing entries were imputed with the metric-wise median. The feature matrix was standardized using global z-scoring, and a single LDA model (n_components=2) was fit on the combined dataset across all windows to obtain LD1/LD2 coordinates in a consistent discriminant space.

## Extended Data Figures

**Extended Data Fig. 1.**
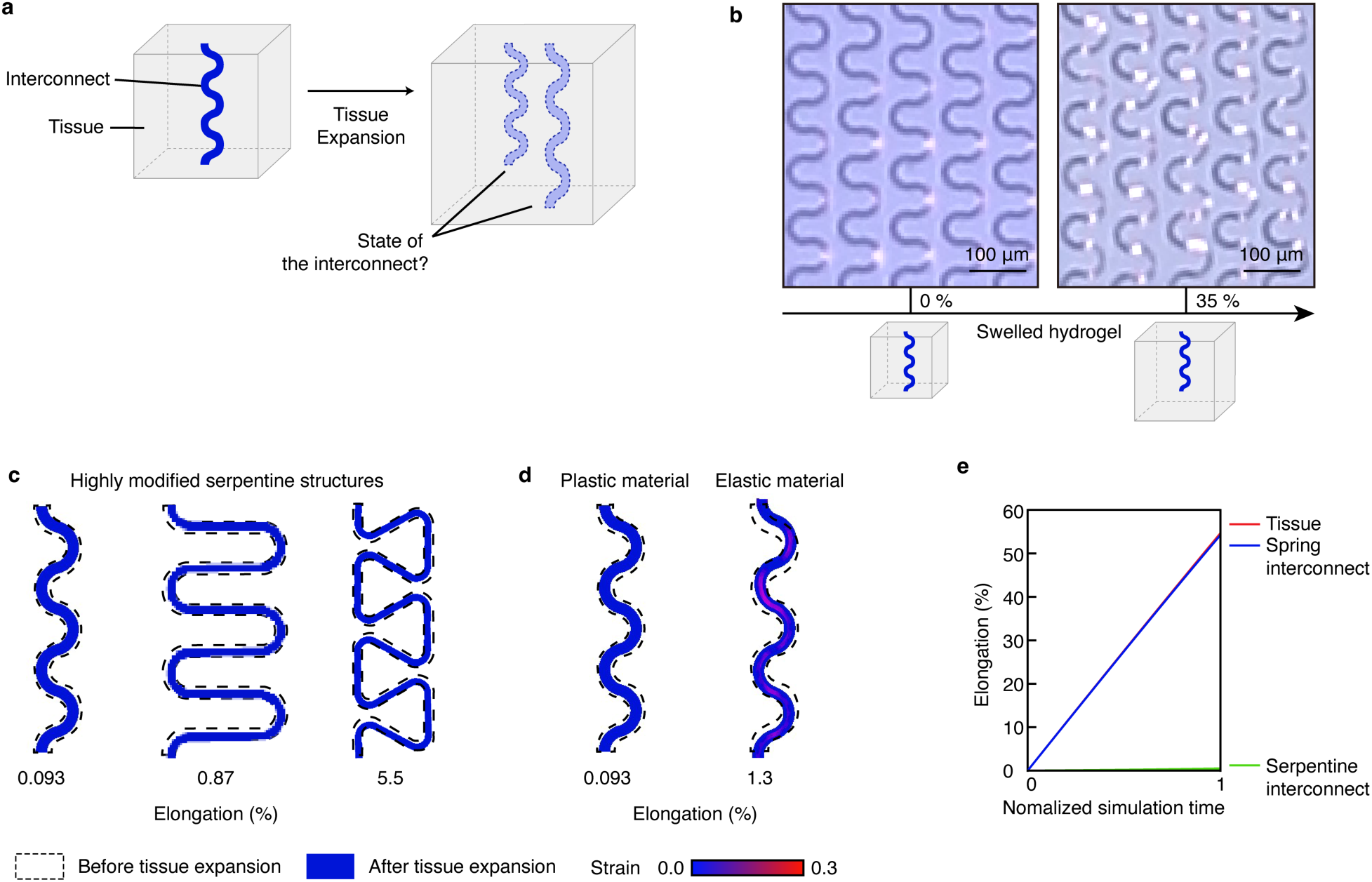
Mechanical simulation and design optimization of interconnects for the developing brain. **a**, A schematic describing the craniocaudal expansion and interconnect relationship. **b**, Microscopy images of serpentine interconnects embedded in a hydrogel brain phantom before and after 35% expansion along the depth direction. **c**, FEA showing three highly stretchable serpentine structure variations under 30% craniocaudal expansion **d**, FEA showing strain distribution in a classic serpentine design fabricated with plastic material (left) or elastic material (right). **e**, Elongation of spring and spiral interconnect within the expanding tissue.

**Extended Data Fig. 2.**
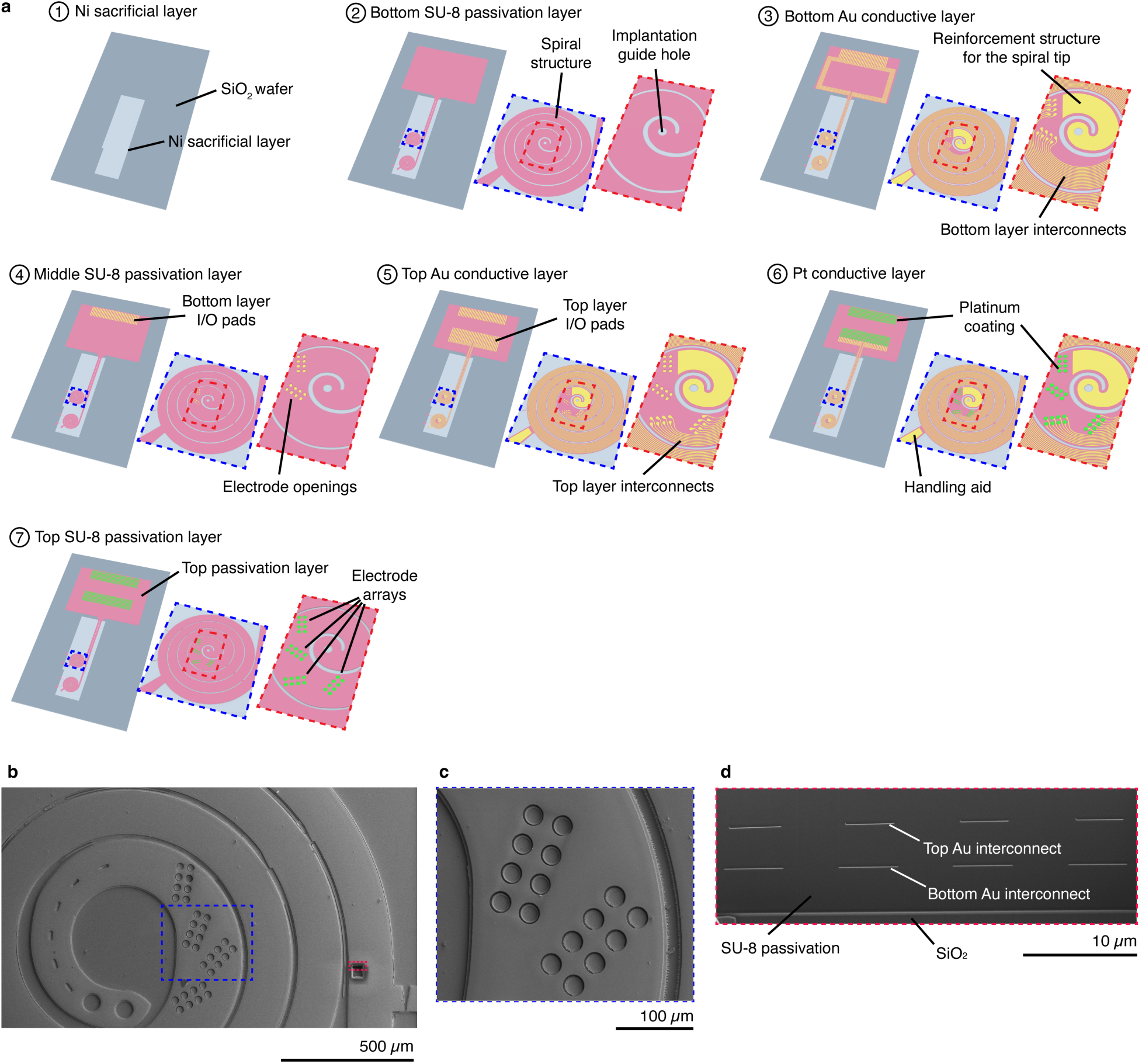
Microfabrication and electrical characterization of spiral-to-spring electronics. **a,** Key microfabrication/assembly steps on a SiO₂ wafer with a Ni sacrificial layer, including formation of the spiral, guide hole, I/O pads, anchors and implantation handle, followed by Pt electrode coating and final passivation. **b**, Scanning electrode microscopy (SEM) image of fabricated spiral-to-spring electronics showing a part of one spiral device. **c**, Close-up SEM image of the blue rectangle region-of-interest from (b) showing eight Pt electrodes. **d**, Cross-section SEM images of the red rectangle region-of-interest from (b) showing the bottom, middle and two layers of Au interconnects (8 channels are shown) embedded in SU-8 passivation layers.

**Extended Data Fig. 3.**
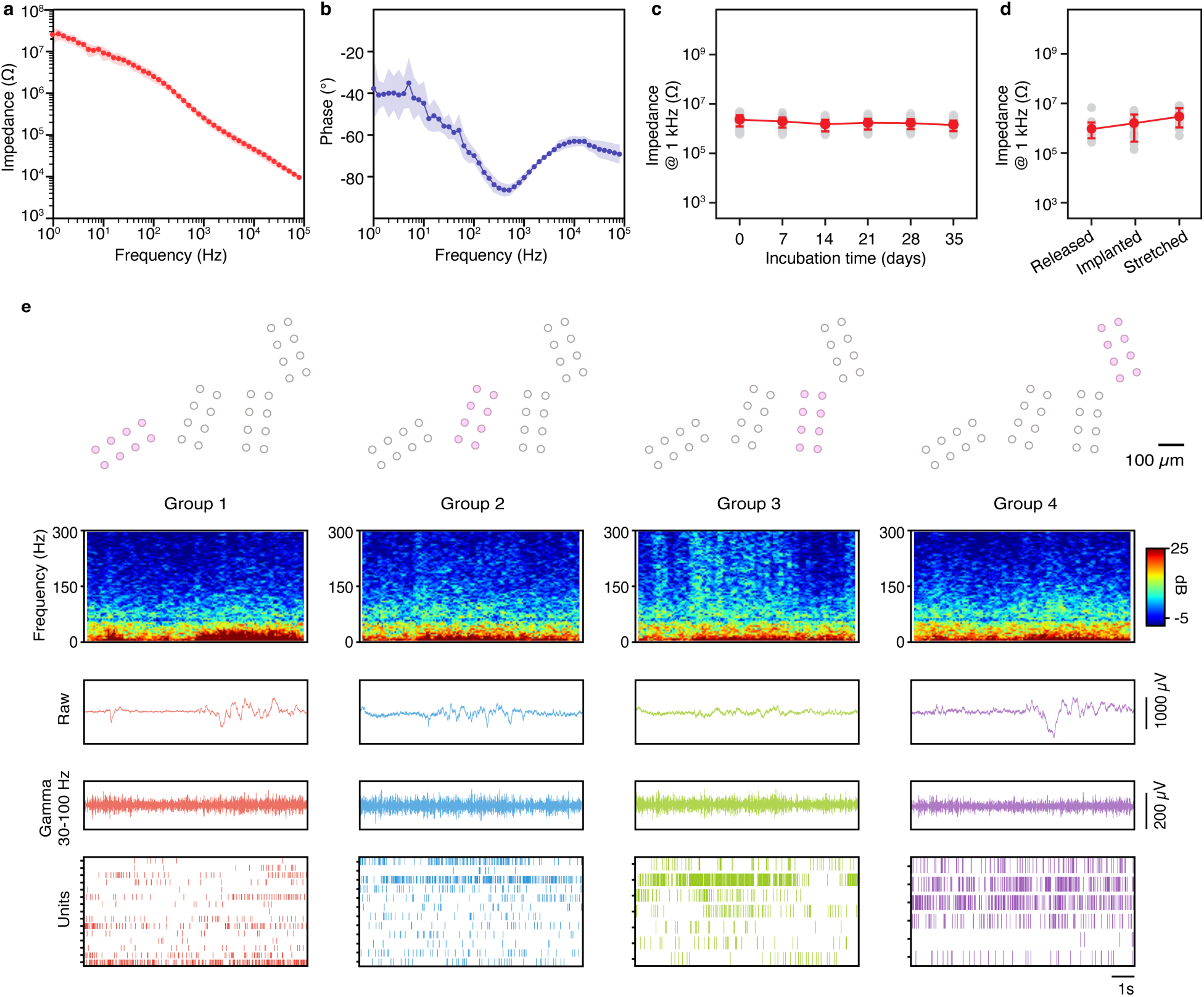
Spiral-to-spring electronics with multi-electrode groups enable wide-band recordings with single unit spikes detected across neonatal development. **a,b**, Amplitude (a) and phase angle (b) of electrochemical impedance spectrum measured from representative device electrodes across frequencies ranging from 0.1 to 100 kHz. **c**, Electrochemical impedance at 1 kHz during phosphate-buffered saline (PBS) incubation remains stable over time. (mean ± s.d.). **d**, Impedance measured after release, implantation into gel and stretching shows no significant degradation (mean ± s.d.). **e**, Top: Schematic of probe electrode configuration with the corresponding group colored with pink, with each group consisting of 8 electrodes. Scale bar: 100 μm. Bottom: Representative recordings from four electrode groups. For each group, local field potential (LFP) spectrogram displaying frequency content from 0.1-300 Hz with power indicated by color scale (top row), raw LFP traces (second row), gamma band traces (third row), and spike raster plots showing sorted unit activity (bottom row).

**Extended Data Fig. 4.**
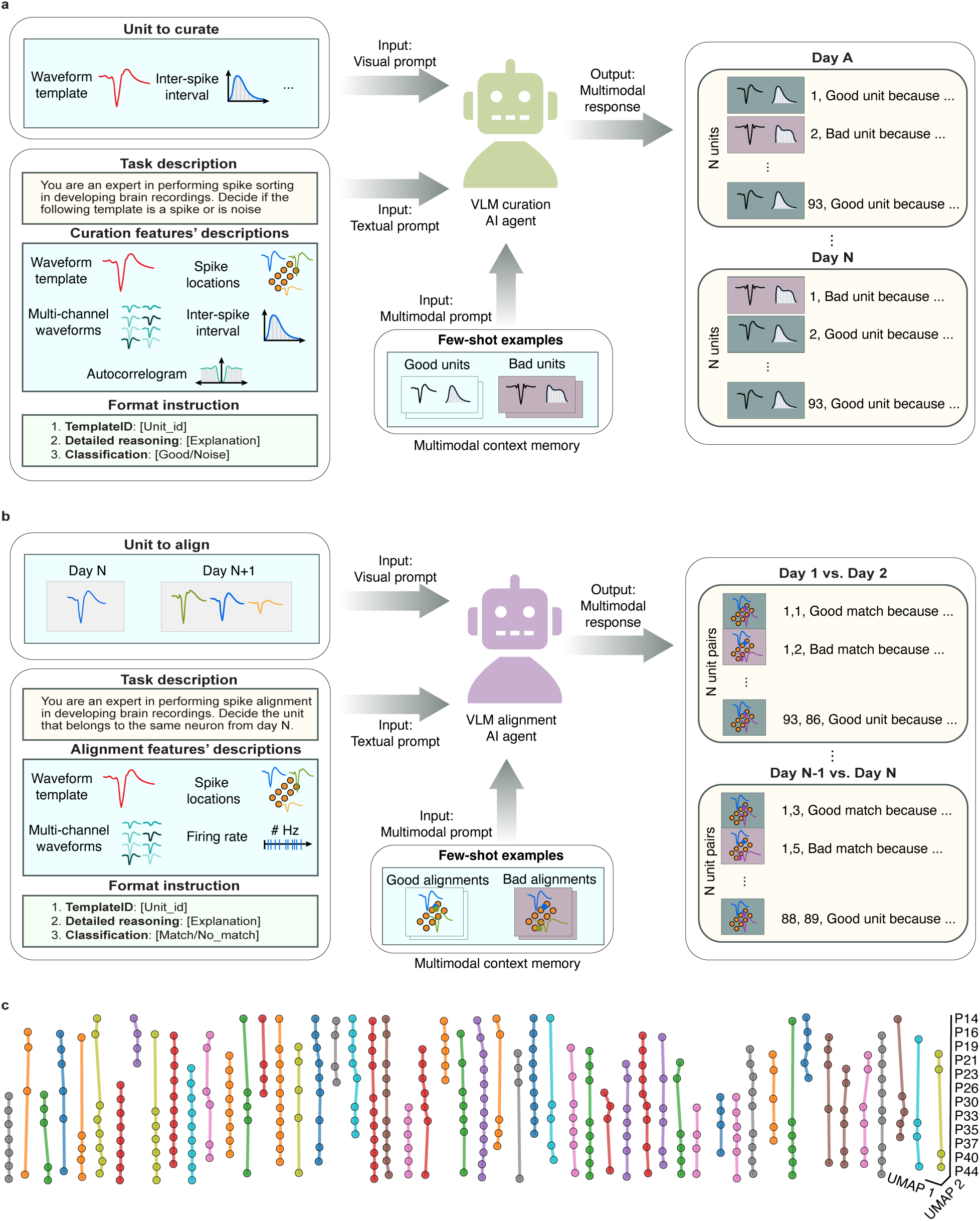
Multimodal agent for spike curation and feature-space priors for a cross-day alignment. **a**, Workflow of the vision-language model (VLM) assistant used to standardize spike curation across sessions. **b**, Schematics describing the workflow of the VLM assistant for cross-day unit alignment. **c**, Uniform manifold approximation and projection (UMAP) of the curated waveform feature vectors across all recorded days in one animal. Points are colored by unit along vertical sequences to illustrate age (P14-P44 as indicated) and provide priors for inter-session alignment of recurring units. Axes denote UMAP dimensions 1 and 2; lines connect consecutive days to emphasize local continuity in feature space.

**Extended Data Fig. 5.**
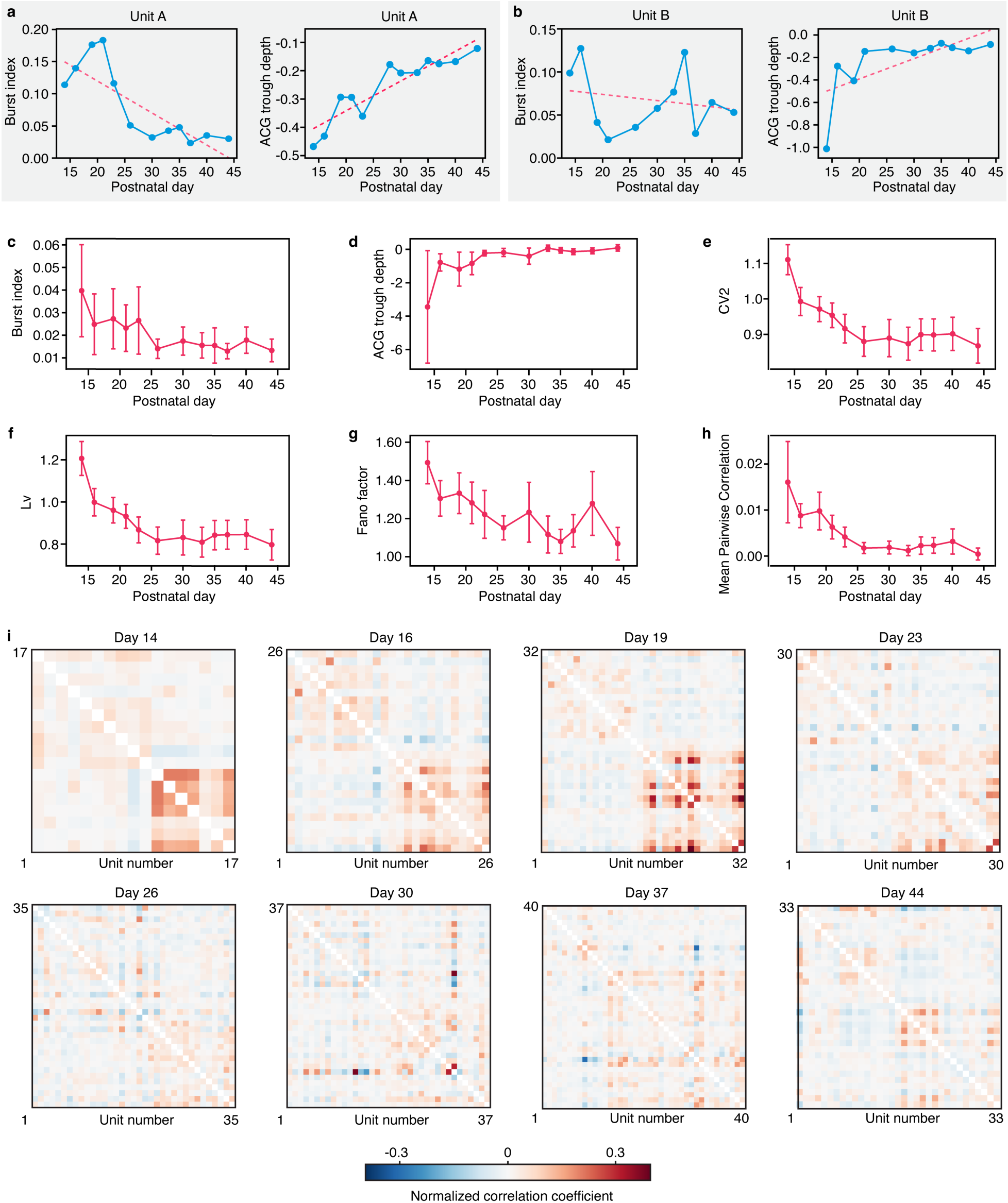
Longitudinal recordings capture hallmarks of cortical maturation. **a,b**, Example developmental trajectories of two representative tracked neurons (Unit A and Unit B) showing burst index (blue solid line, left y-axis) and autocorrelogram (ACG) trough depth (red dashed line, right y-axis) across postnatal development. **c-h**, Population-level developmental trajectories (mean ± SEM) of key electrophysiological metrics across postnatal age (P12-P45). The metrics shown are burst index (c), ACG trough depth (d), local coefficient of variation of ISIs (CV2) (e), Local variation of ISIs (Lv) (f), Fano Factor (g), and mean pairwise correlation (h), **i**, Representative pairwise Pearson correlation matrices at eight developmental time points (P14, P16, P19, P23, P26, P30, P37, P44) from a representative animal. Color bar shows normalized correlation coefficients from −1 to 1 and diagonals are unity (white). Axes denote unit indices.

**Extended Data Fig. 6.**
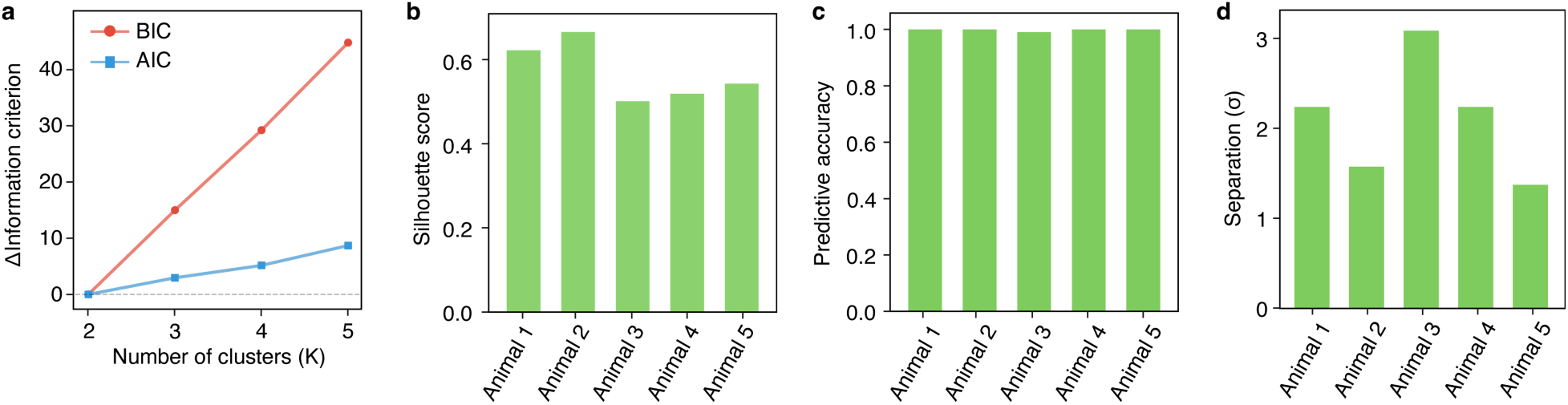
Two-component Gaussian mixture model achieves robust clustering performance across all animals. **a**, Model selection analysis comparing information criteria across different numbers of clusters (K=2-5). **b**, Silhouette scores for per-animal GMM clustering. **c**, Cross-validated area under the receiver operating characteristic curve (CV AUROC) for cluster assignment prediction. **d**, Component separation between the two GMM cluster means, expressed in units of pooled standard deviations (σ), for each animal.

**Extended Data Fig. 7.**
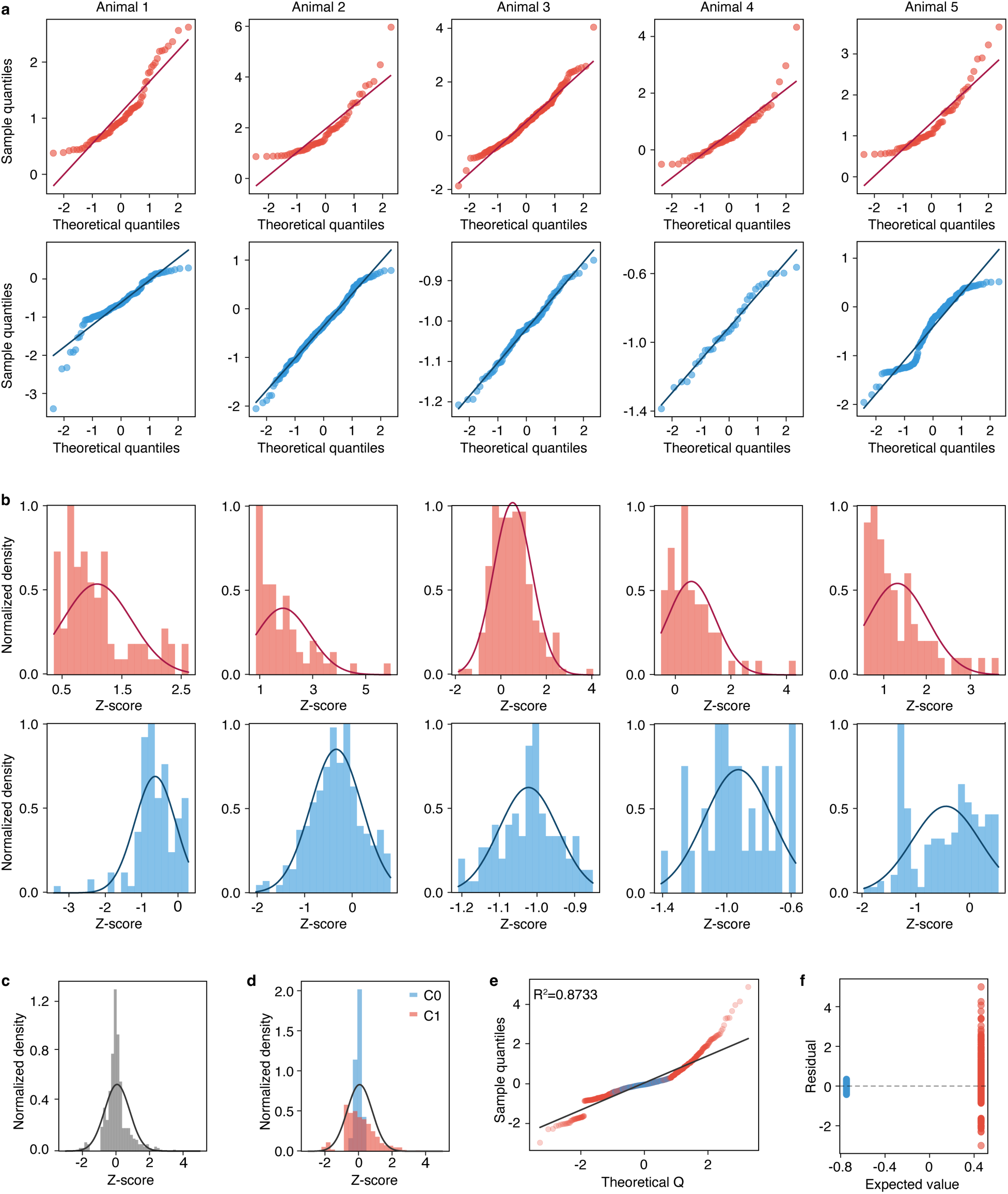
Population coupling values within GMM-defined clusters approximate Gaussian distributions, validating the mixture model assumption. **a**, Quantile-quantile (Q-Q) plots comparing the distribution of population coupling z-scores within each cluster against theoretical normal distributions, shown separately for each animal. Top row: high-population coupling cluster (C1, red); bottom row: low-population coupling cluster (C0, blue). Points falling along the diagonal reference line indicate agreement with normality. **b**, Histograms of population coupling z-scores within each cluster overlaid with fitted Gaussian curves. Top row: C1; bottom row: C0. **c**, Distribution of residuals (observed − expected values based on cluster mean assignment) pooled across all neurons, with fitted Gaussian overlay (gray). **d**, Residual distributions shown separately by cluster. **e**, Q-Q plot of pooled residuals colored by cluster assignment. **f**, Residuals plotted against expected values (cluster means). Residuals are centered around zero for both clusters.

**Extended Data Fig. 8.**
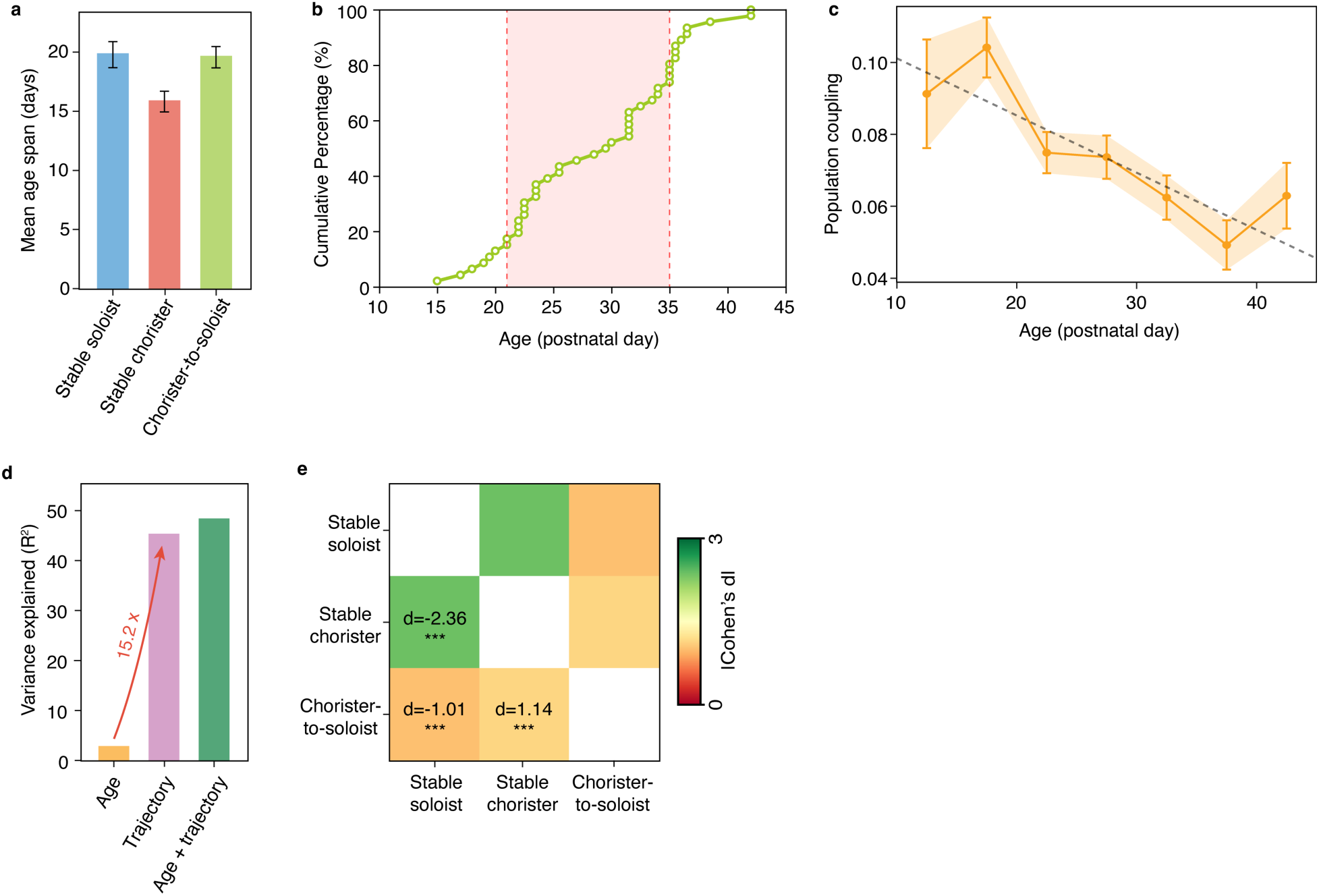
Longitudinal mapping reveals three distinct developmental trajectories of population coupling that are obscured by cross-sectional analysis. **a**, Mean age span (days) of longitudinal mapping for each trajectory type: stable soloist, stable chorister, chorister-to-soloist. **b**, Cumulative distribution of transition timing from chorister-level population coupling to soloist-level population coupling for chorister-to-soloist trajectory neurons. Shaded region indicates critical period. **c**, Cross-sectional analysis of population coupling across postnatal age, pooling all neurons regardless of trajectory (gray line, mean ± SEM). **d**, Variance decomposition comparing the explanatory power of different predictors. The red arrow indicates the fold-increase in explained variance. **e**, Pairwise effect size matrix with statistical significance. All comparisons are highly significant (***p<0.001, two-sided independent t-tests). Color scale represents absolute effect size magnitude.

**Extended Data Fig. 9.**
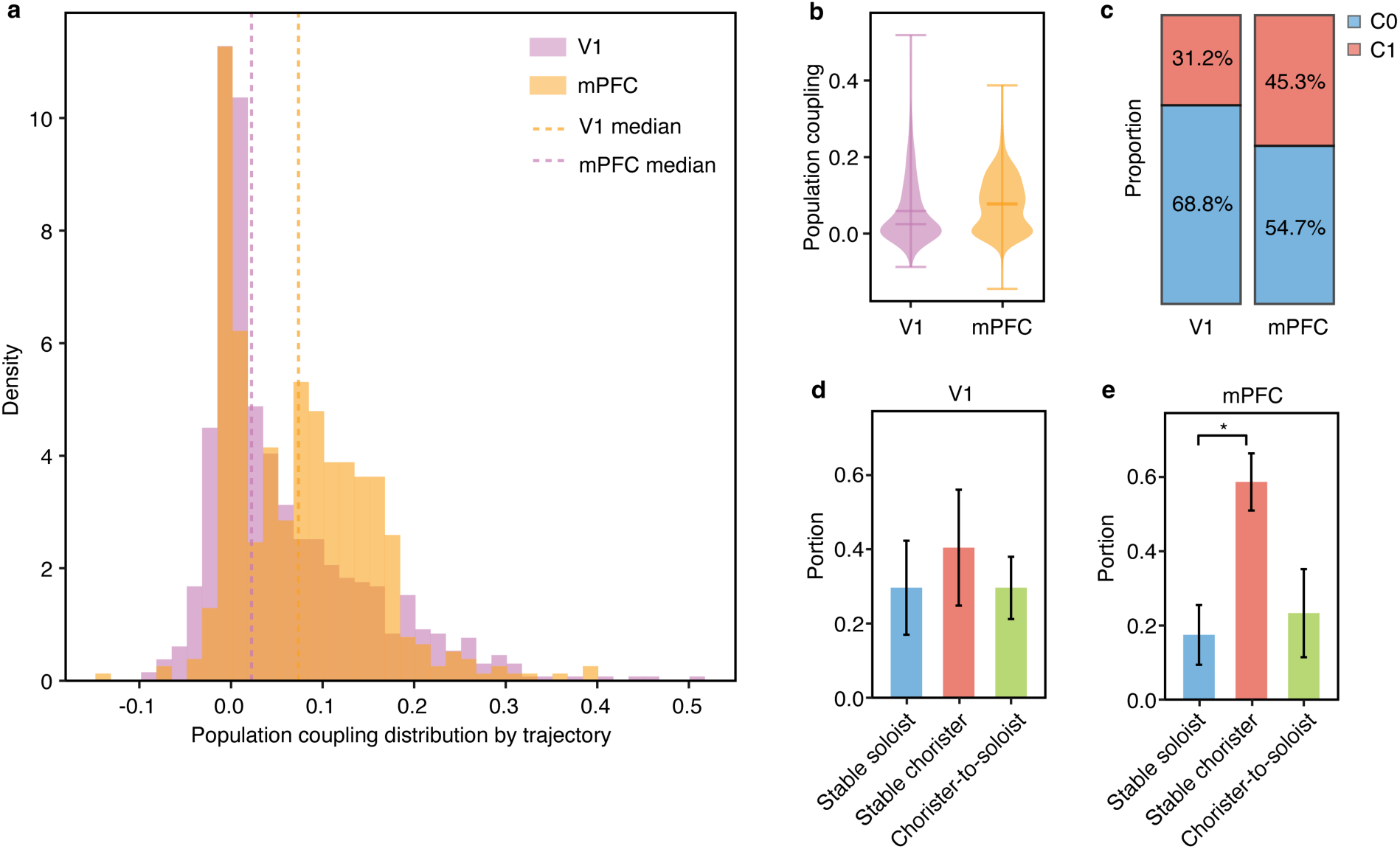
Visual cortex and medial prefrontal cortex exhibit distinct population coupling distributions and developmental trajectory compositions. **a**, Overlapping histograms showing the distribution of population coupling values in visual cortex (V1, purple) and medial prefrontal cortex (mPFC, orange). Dashed vertical lines indicate median values for each region. **b**, Violin plots comparing population coupling distributions between V1 and mPFC, showing the full distribution shape, mean, and median for each region. **c**, Stacked bar chart showing the proportion of neurons assigned to the low-population coupling cluster (C0, blue) versus high-population coupling cluster (C1, red) in each brain region. **d,e**, Bar plots showing trajectory proportions by brain region (d) V1, visual cortex and (e) mPFC, medial prefrontal cortex.

**Extended Data Fig. 10.**
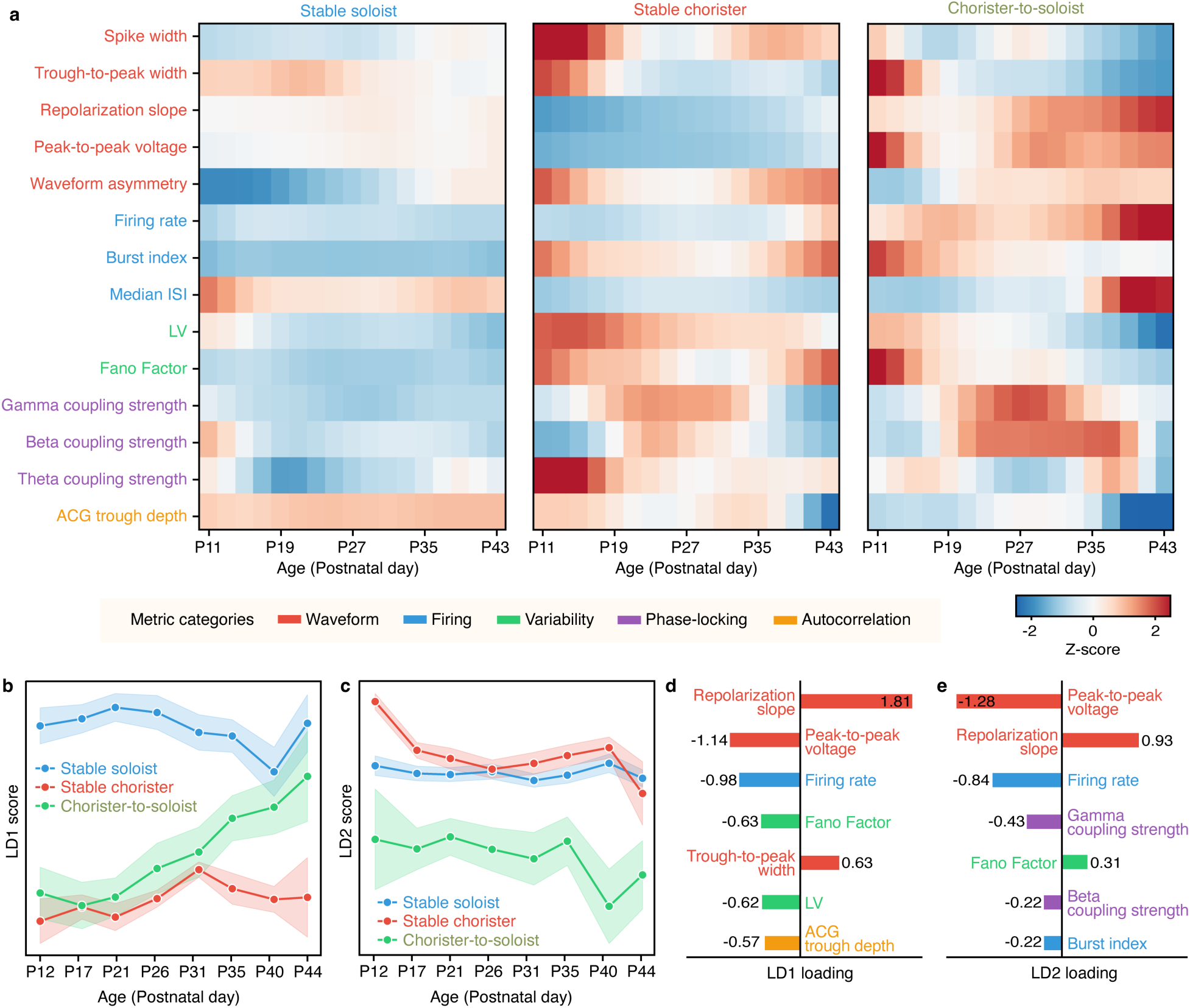
Developmental trajectories of other intrinsic neuronal properties differ in stable choristers, chorister-to-soloists, and stable soloists across postnatal maturation. **a**, Heatmap showing developmental changes in 14 independent electrophysiological metrics across three trajectory types (stable soloist, stable chorister, chorister-to-soloist) from postnatal day 11 (P11) to P43. Values are Z-score normalized within each metric (row) across all trajectory-age combinations, then Gaussian smoothed (σ=1.5). Blue indicates values below the mean; red indicates values above the mean. Metrics are grouped by functional category: waveform properties (spike width through waveform asymmetry), firing properties (firing rate through median ISI), spike timing variability (LV, Fano Factor), oscillatory coupling strength (gamma through theta), and autocorrelation feature (ACG trough depth). **b,c**, Developmental trajectory of primary discriminant axis (LD1) (b) and secondary discriminant axis (LD2) (c) from linear discriminant analysis (LDA). Shaded envelops indicates within-bin variability. **d,e**, Metric contributions to LD1 (d) and LD2 (e).

